# Transgenerational Epigenetic Inheritance of MHC Class I Gene Expression is Regulated by the CCAAT Promoter Element

**DOI:** 10.1101/2023.04.13.536772

**Authors:** Jocelyn D. Weissman, Aparna Kotekar, Zohar Barbash, Jie Mu, Dinah S. Singer

**Affiliations:** Experimental Immunology Branch, Center for Cancer Research, National Cancer Institute, NIH, Bethesda, MD 20892

**Author notes:** Current Address: NIH Center for Human Immunology, Inflammation, and Autoimmunity (CHI), National Institute of Allergy and Infectious Diseases, NIH, Bethesda, MD 20892. Current address: FORE biotherapeutics, Philadelphia, PA. Co-first authors (Jocelyn D. Weissman, Aparna Kotekar). Corresponding author Dinah S. Singer, Bldg 10, Room 4B-36 NIH, Bethesda, MD 20892.

## Abstract

Transgenerational epigenetic inheritance is defined as the transmission of traits or gene expression patterns across multiple generations that do not derive from DNA alterations. The effect of multiple stress factors or metabolic changes resulting in such inheritance have been documented in plants, worms and flies and mammals. The molecular basis for epigenetic inheritance has been linked to histone and DNA modifications and non-coding RNA. In this study, we show that mutation of a promoter element, the CCAAT box, disrupts stable expression of an MHC Class I transgene, resulting in variegated expression among progeny for at least 4 generations in multiple independently derived transgenic lines. Histone modifications and RNA polII binding correlate with expression, whereas DNA methylation and nucleosome occupancy do not. Mutation of the CCAAT box abrogates NF-Y binding and results in changes to CTCF binding and DNA looping patterns across the gene that correlate with expression status from one generation to the next. These studies identify the CCAAT promoter element as a regulator of stable transgenerational epigenetic inheritance. Considering that the CCAAT box is present in 30% of eukaryotic promoters, this study could provide important insights into how fidelity of gene expression patterns is maintained through multiple generations.

## INTRODUCTION

Transgenerational epigenetic inheritance (TEI) is the process by which novel phenotypes are transmitted from one generation to the next through epigenetic, not DNA sequence, changes (1,2). It has been widely observed and documented in plants. The number of examples of TEI in mammals is still limited but increasing (3). TEI is induced by environmental factors such as toxic exposure, stress, starvation or obesity, that cause phenotypic changes in multiple subsequent generations without an accompanying change in DNA sequence.

Although the precise mechanisms underlying TEI are still not known, a number of epigenetic associations have been established in both plants and mammals. Among the documented links with epigenetic inheritance are changes in DNA methylation, histone modifications and small RNAs (1,2,4). It is well established that plants maintain epigenetic inheritance through DNA methylation which is stably transmitted through generations thus affording a mechanism for the formation of epialleles in the progeny (1). In contrast to plants, mammalian DNA undergoes complete demethylation in the germline with subsequent restoration of parent-specific imprints which lead to parent-of-origin phenotypic inheritance. In mammals, transposons, which are often resistant to demethylation, are among the best characterized examples of epigenetic inheritance (5). Histone modifications have been shown to determine epigenetic inheritance independent of DNA methylation in C. elegans and Drosophila, where H3K27me3 marks are transmitted to progeny. In the mouse, H3K27me3-dependent imprinting has been observed where it is established during oogenesis but only persists during early embryogenesis (6). Finally, germline specific noncoding RNAs have been associated with establishing epigenetic inheritance following various exposures (7). Evidence to date for TEI in mammals is largely limited to transposons, transgenes inserted next to transposons or intracisternal A particles. Absent from any of these studies is evidence that DNA sequences are involved in establishing or preventing TEI.

Major histocompatibility complex class I genes are ubiquitously expressed, but their level of expression varies among tissues and is determined by both tissue-specific and hormonal signals. The MHC class I promoter consists of elements homologous to the canonical elements CCAAT, TATAA, Sp1 binding site (Sp1BS), and Initiator (Inr). The CCAAT element is completely conserved among MHC class I genes within a family and among mammalian species (8). Consistent with the CCAAT box playing a critical role in transcriptional regulation, CCAAT box mutations are causative for a number of human diseases, including hemoglobinopathies. Although a number of transcription factors bind to the CCAAT box, the most common is the trimeric complex, NF-Y, which regulates expression of the major histocompatibility (MHC) class I and class II genes, as well as globin and albumin genes (9–12). Mutations in the CCAAT box leads to loss of NF-Y binding and reduced expression of the associated gene.

As we have reported previously, a transgene spanning the MHC class I gene, PD1, was expressed in mice with the same tissue-specific and cytokine-dependent patterns as the endogenous class I genes (13,14). Surprisingly, as assessed by individually mutating each of the promoter elements in the context of the transgene, none were essential for class I expression. In particular, we found that the CCAAT box regulates constitutive expression of the MHC class I gene in a tissue-specific manner. Thus, mutation of the CCAAT box in the context of the MHC class I promoter does not affect constitutive expression in the spleen. However, it leads to significantly higher expression in kidney and brain indicating it functions as a repressor in non-lymphoid tissues (14). These studies identified the CCAAT box as a tissue specific regulator of MHC class I transcription, functioning as an activator or repressor element in a context-specific fashion.

In the present studies, we extend the characterization of the CCAAT box and report that it also functions to maintain stable MHC class I expression across generations and abrogate epigenetic transgenerational patterns of expression. Among multiple independent CCAAT mutant (CCAATm) transgene founders, some founders expressed the transgene whereas others did not. However, in subsequent generations, progeny of non-expresser founders expressed the transgene. Those progeny, in turn, gave rise to transgenic progeny that were non-expressers. This generational alternation of expression occurred over multiple generations until stabilizing after a few generations, such that transgenic lines derived from a common founder displayed distinct patterns of expression. In contrast to the wild type CCAAT box, NF-Y does not bind the mutated CCAAT box. The CCAATm transgene expressers are distinguished from the non-expressers by their patterns of histone modification, Pol II and CTCF binding and DNA looping. These studies demonstrate that the conserved MHC class I CCAAT box functions both as a transcriptional regulator and to maintain stable expression; mutation of the CCAAT box leads to variegated expression between generations.

### Material and methods

#### Mice

C57BL/10 mice homozygous for the MHC class I transgene PD1 (CCAATwt) were generated as described previously [1]. The CCAATwt transgene contains a 1 Kb regulatory region upstream of the TSS, the entire coding region and 650bp immediately following the polyadenylation site that contains the 3’ boundary element (15). Experiments were performed on CCAATm expresser and non-expresser lines derived from at least two independent founders [3]. Peripheral blood lymphocytes (PBL) were analyzed by flow cytometry of cells stained with anti-class I antibody. All animal procedures reported in this study that were performed by NCI-CCR affiliated staff were approved by the NCI Animal Care and Use Committee (ACUC) and in accordance with federal regulatory requirements and standards. All components of the intramural NIH ACU program are accredited by AAALAC International.

#### IFN treatment

Mice were injected intraperitoneally with 50kU of mouse IFNγ (CalBiochem) or an equal volume of PBS + 0.1%BSA. Tissues were harvested 24 hours post-injection. Levels of PD1 and endogenous H2-K^b^ RNA were assessed by RT real time PCR of RNA extracted from tissues. The experiment was performed twice each with three mice that were analyzed independently.

#### Flow cytometry

FACS was carried out using antibodies for PD1 surface expression (PT85, VRB) as described before (16). FACS results were analyzed using FlowJo [BD].

#### Immunofluorescence

Immunofluorescence staining was performed according to standard protocol, using the PT85 antibody that was used for FACS staining. Slides were counterstained with DAPI.

#### Electrophoretic Mobility Shift Assay (EMSA)

NF-Y EMSA was done as described previously (17) with modifications mentioned below. DNA fragments used in the gel shift assay were the PD1 fragment from 1011-1060 and the corresponding CCAAT mutant fragment (IDT). DNA fragments were end labeled with [32P]ATP using T4-Polynucleotide kinase (NEB). EMSAs were done with 3.6ug of HeLa nuclear extract and 1.5-fmol radiolabeled probe in binding buffer (20mMTris-HCl pH 7.5, 50 mM NaCl, 5 mM MgCl2, 0.5 mM EDTA, 6.5% glycerol, 2.5 mM DTT, and 0.1 mg/ml BSA) (18) and 2ug poly(dA-dT). Reactions were incubated at 30^°^C for 30’. For cold competition and antibody supershifts, cell extracts were preincubated with the competitor or antibody for 30 min on ice before addition of radiolabeled probe. Competitor double-stranded oligonucleotides were added at a 1000-fold molar excess. Antibodies used were NF-YA Antibody (G-2): sc-17753, and NF-YB Antibody (G-2): sc-376546 (both from Santacruz). The reactions were loaded on 4% polyacrylamide gels and run in 0.5X TBE at 200 V for 2 hours.

#### MNase hypersensitivity and Nucleosome Occupancy

Nuclei were prepared from spleen cells from CCAATwt and CCAATm expresser and non-expresser mice as described previously (19). These nuclei were used for MNase digestions as described before (19). Briefly, 15×10˄6 nuclei were digested for 5’ at 37°C with 0 or 3U of micrococcal nuclease in 100ul of MNase digestion buffer with 1mM CaCl2. Nucleosome occupancy was assessed across the PD1 gene (–614, –30, Exon 1 and Exon 5; primers below) by real-time PCR, measured as amplification from 100ng digested DNA (3U MNase) normalized to undigested (0U) DNA.

Additionally, nuclei were digested with high MNase concentrations 50U and 100U for 5’ and 15’ at 37°C to assess accessibility at the PD1 promoter (CCAAT box) by amplifying DNA from the digested soluble chromatin fraction versus the digestion resistant pellet fraction.

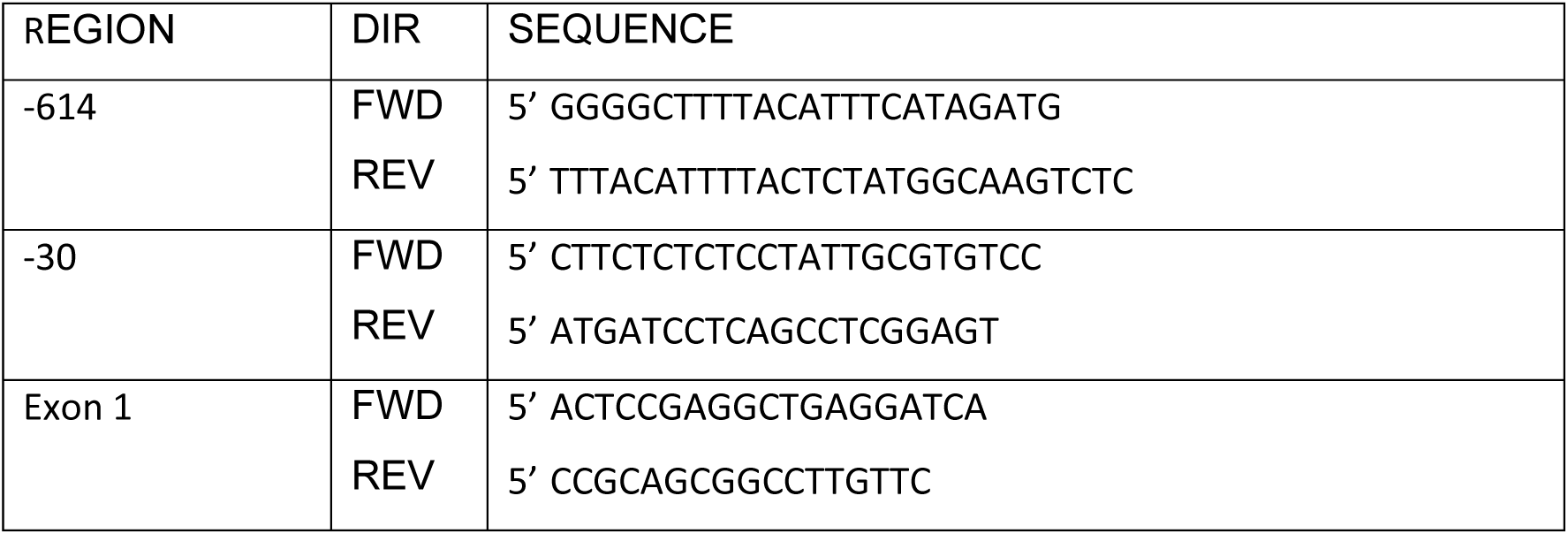
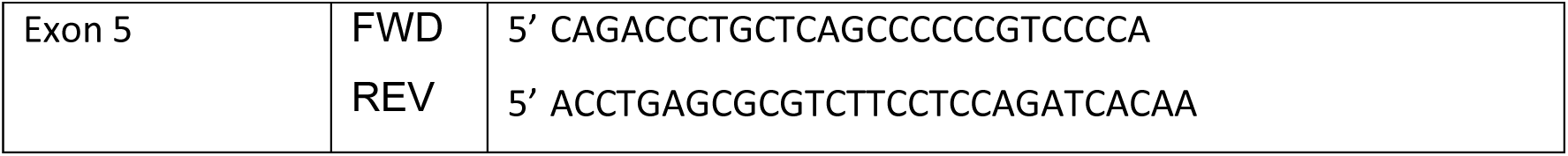

#### Real-time RT-PCR

RNA was prepared using RNeasy kit (Qiagen) according to manufacturer instructions. Each sample was subjected to DNase treatment on column using RNAse free DNase (Qiagen) per manufacturer’s instructions. cDNA was synthesized using oligo dT or random primers and the Superscript III kit according to manufacturer’s instructions (ThermoFisher). Real time PCR was performed as described (19) using ABI7900 with SYBR PCR Master Mix (Life technologies). For tissue RNA expression, the calculations used the standard curve method, and normalized to the level of 18S in the tissues. All results reported are the average of 2-3 independent experiments in lines derived from different founders, to exclude possible insertion position effects. Endogenous MHC class I levels were determined the same way using published primers (20).

#### Assessment of upstream transcription

Tissues were harvested from CCAATwt and CCAATm expresser and non-expresser mice and stored at –80 degrees C. RNA was prepared by using TRIzol (Invitrogen/Thermo Fisher Scientific). DNA contamination was eliminated using TURBO™ DNA-free kit (Invitrogen/Thermo Fisher Scientific). The RNA was then used for cDNA preparation using AffinityScript Multiple Temperature cDNA Synthesis Kit (Agilent) with random primers, oligodT primers or gene-specific primers (Integrated DNA Technologies).

Gene-specific cDNA primers used were:

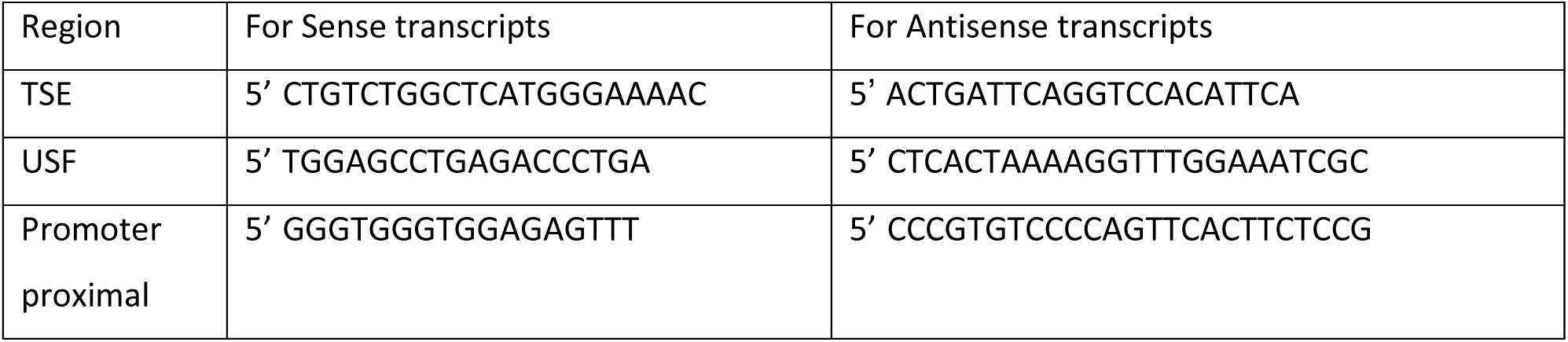

cDNAs were then amplified using SYBR green PCR master mix (Applied Biosystems,). Primers used for real time RT-PCR were:

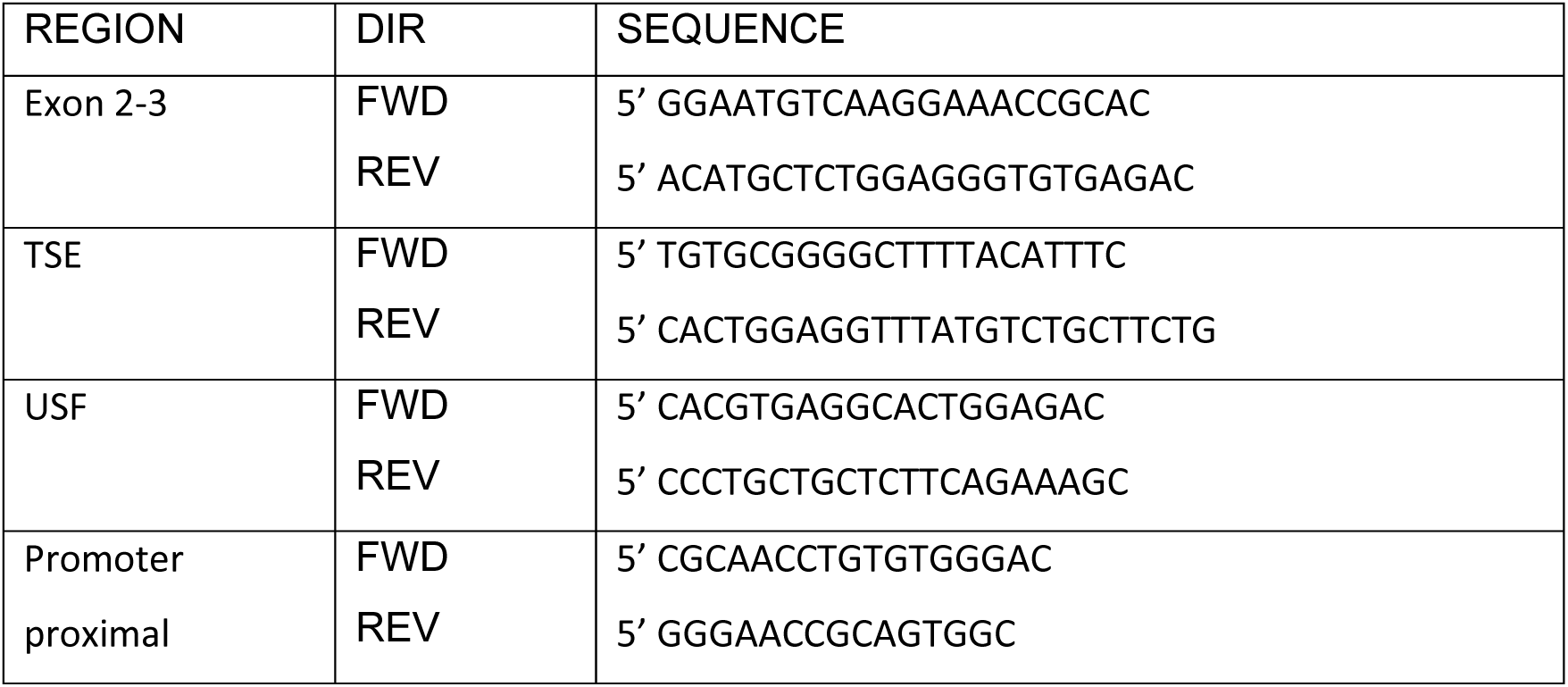

Primers for 18S RNA (AM1716, Ambion/Applied Biosystems) were used for amplifying the internal control. Results were normalized for 18S and copy number.

#### NF-Y siRNA treatment and effect on PD1 expression

93B2 cells (Mouse L-cells stably transfected with the PD1 gene) were cultured for 24-hours in 12-well plates and then transfected in duplicate with NF-Y siRNA (Mm_Nfya_3 FlexiTube siRNA, or Mm_Nfya_2 FlexiTube siRNA, or Mm_Nfyb_4 FlexiTube siRNA; Qiagen) using Lipofectamine RNAiMax Reagent (Thermofisher). GAPDH siRNA (Silencer™ Select GAPDH Positive Control siRNA; Thermofisher) and a non-targeting siRNA pool (ON-TARGETplus Non-targeting Control Pool; Horizon/Perkin Elmer) were used as controls. The cells were harvested at 24, 48 and 72 hours. RNA was extracted using RNAeasy plus mini kit. cDNA was prepared using AffinityScript Multiple Temperature cDNA Synthesis Kit (Agilent) with random primers, and amplified using SYBR green PCR master mix (Applied Biosystems, Foster City, CA).

siRNA knockdown was confirmed by RT-PCR using NF-YA, NF-YB and GAPDH primers, and effect of NF-Y knockdown on PD1 expression was assessed by RT-PCR using PD1 primers. Primers for 18S RNA (AM1716, Ambion/Applied Biosystems) were used for amplifying the internal control.

Primers used for real time RT-PCR were:

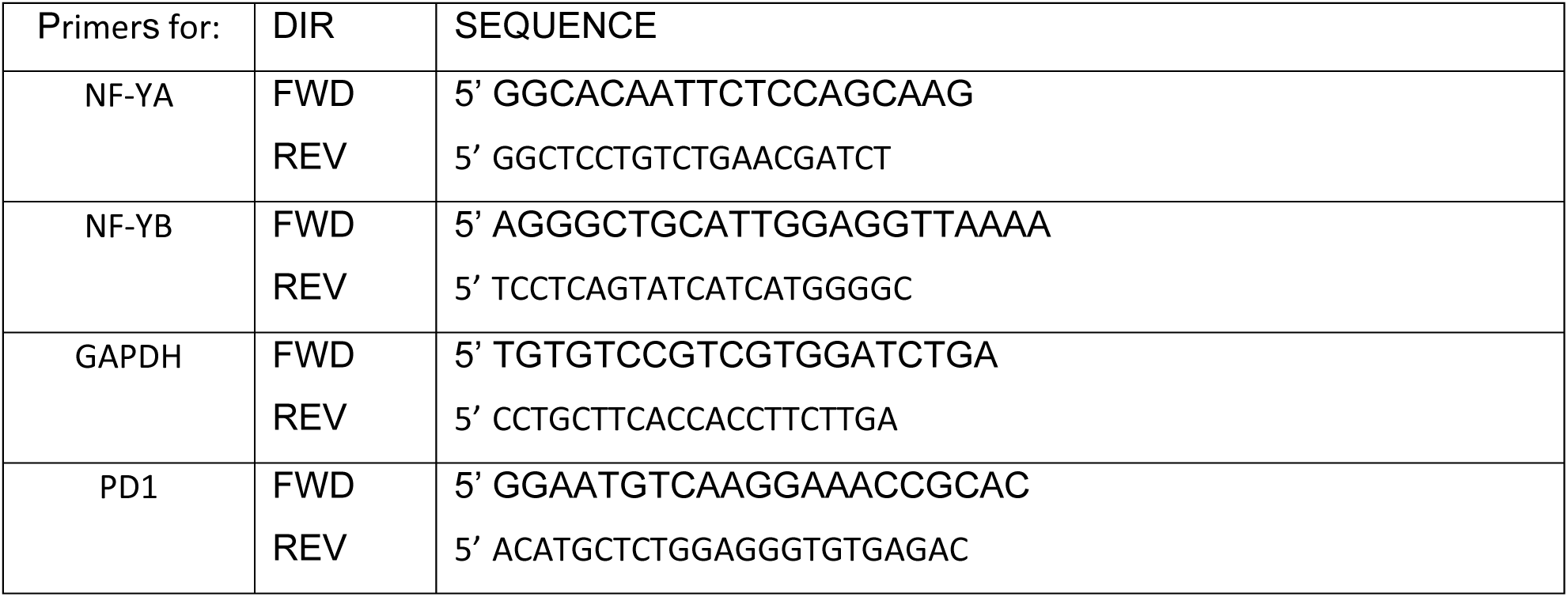

#### Chromatin Immunoprecipitation (ChIP)

Spleens from 2 transgenic mice were pooled for each experiment. The lysis of the tissues was performed by the MAGNA ChIP Kit for tissues (Millipore). ChIPs were performed according to manufacturer instructions. All experiments were repeated 3 times. Antibodies used: anti Pol-II (Active Motif, Santa Cruz), anti NF-YB (Santa Cruz Biotechnology), anti AcH3 (Abcam), anti Smc1 (Millipore), anti CTCF (Millipore, Santa Cruz), anti H3K4me3, H3K9me3 (Upstate)

#### Quantitation of ChIP results

DNA immunoprecipitated in ChIP reactions was analyzed by real time PCR using the primers described below. (Real-time PCR, rather than ChIP-seq, was used since the analyses focused a transgene.) Quantitative real-time PCR was performed using ABI7900 with SYBR green real time PCR kit (Applied Biosystems). Results were calculated as percentage bound/total input DNA, relative to IgG control.

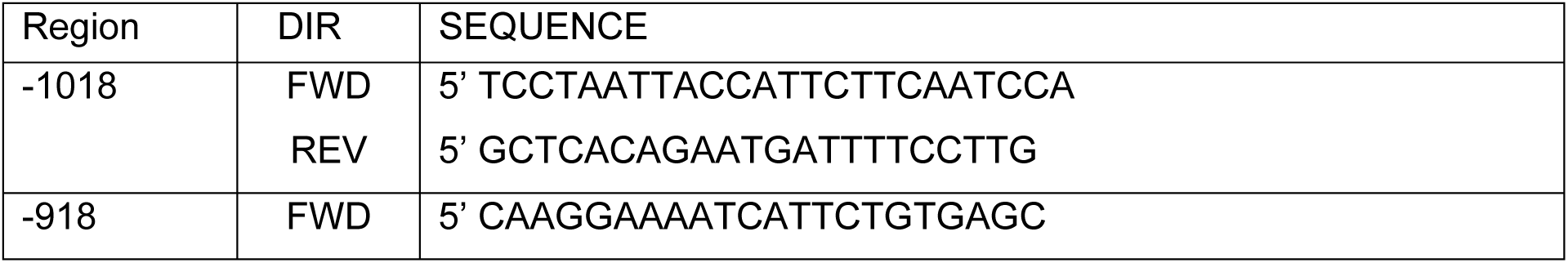
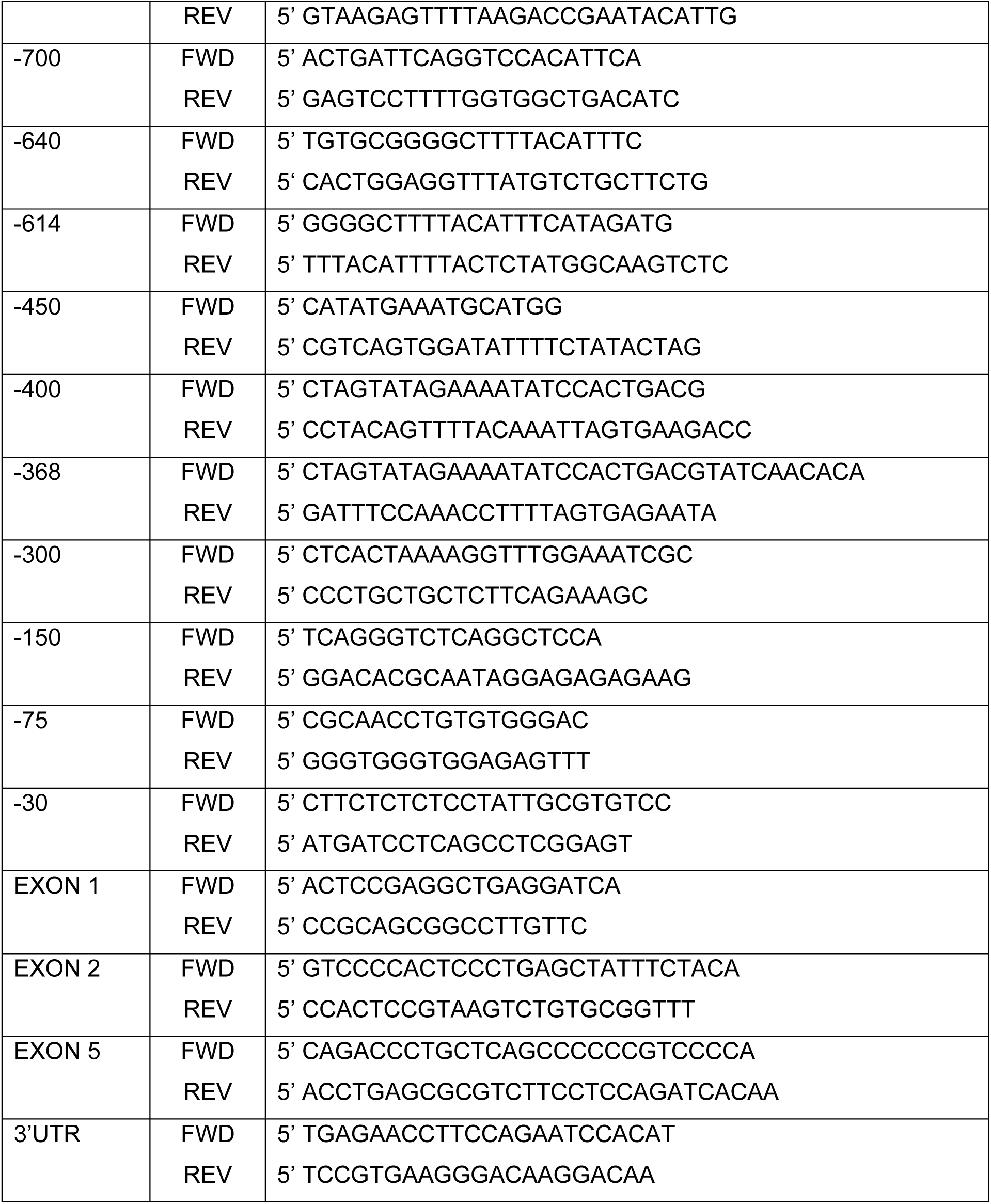

#### Spleen Chromosome conformation capture 3C Analysis

3C analysis was performed as described (21,22), using Nla III [NEB special order] as the restriction enzyme. Single cell suspensions of ACK treated spleen cells were crosslinked at RT for 5 minutes with 1% formaldehyde in HBSS/2%FBS, then quenched with glycine to 0.125M for 15min on ice. Washed crosslink cells were lysed by swelling cold and harvested by rapid centrifugation to obtain nuclei. 20E6 cell equivalents were used for each experimental point. A non-digested sample was kept. 4000 Units of NlaIII [custom concentration 50,000Units/ml] was added and samples were incubated rotating overnight at 37⁰C. The following morning, enzyme was inactivated with SDS and ligations [T4 DNA ligase, NEB] of varied concentrations were set up for 4 hours at 16⁰C. Samples were RNAse treated, phenol and IAC extracted, and precipitated. PCRs were performed to determine ligated sites. Non-digested, and digested non-ligated samples were used as controls. PCR analysis of 3C experiments were performed using primers in table below.

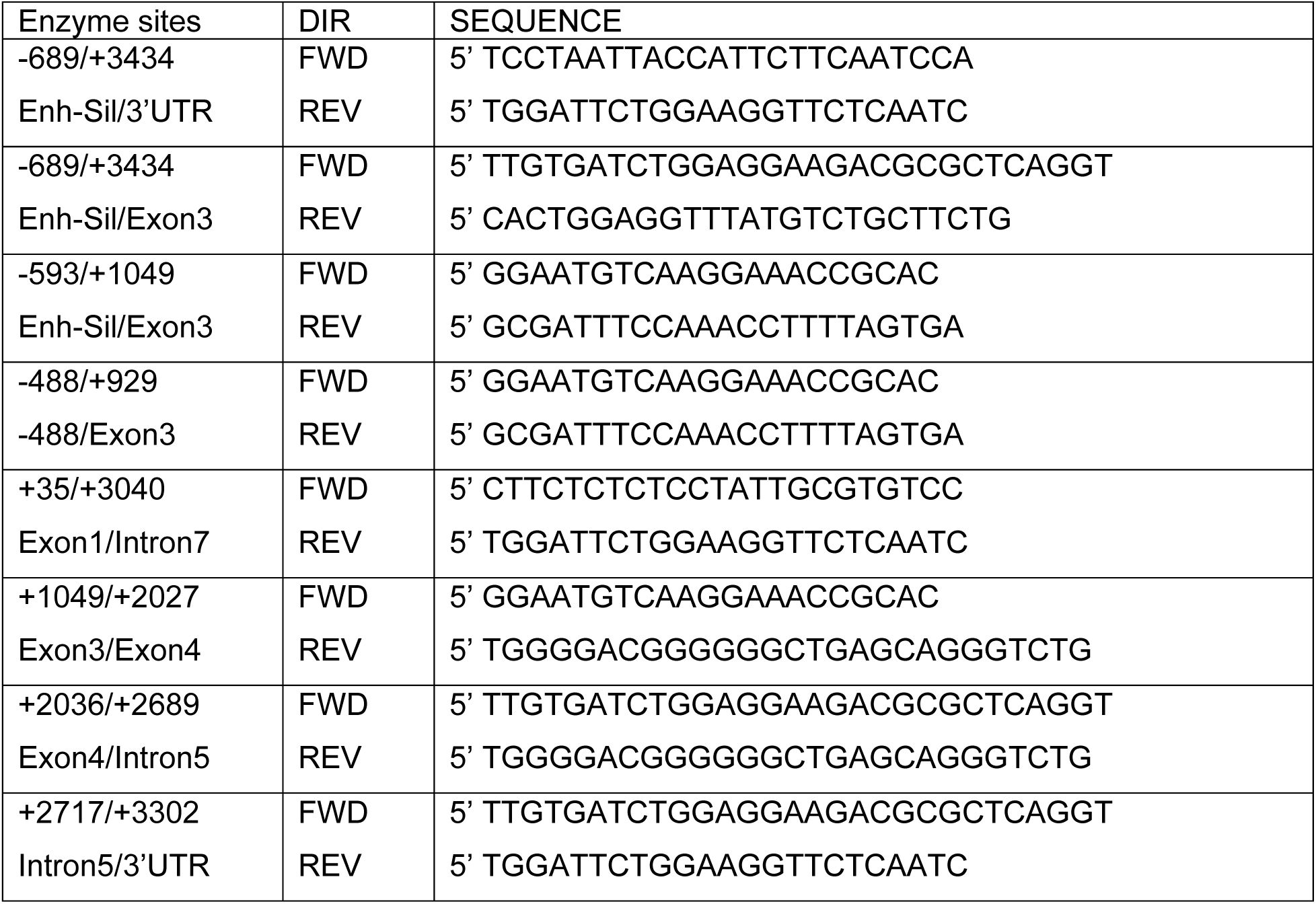

#### Methylation assay

Methylation of genomic DNA was examined using methylation sensitive enzymes Aci 1 or Tai 1. 2 ugms of genomic DNA from spleens of CCAATwt and CCAATm mutant mice, were digested with 20 units of either enzyme overnight. PCRs were then performed using oligos flanking the enzymatic sites on the digested DNA and compared to undigested.

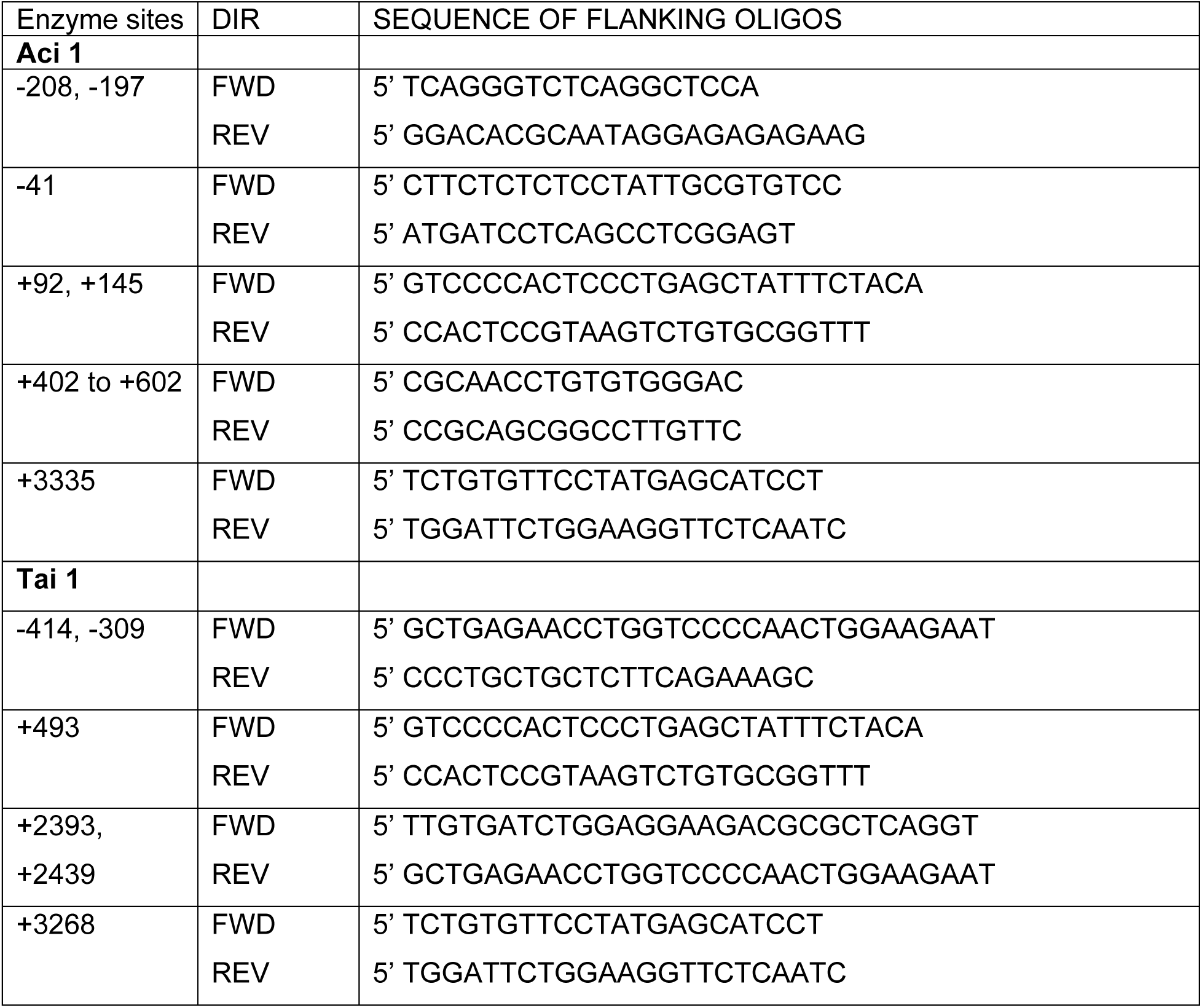

## Results

### Mutation of the CCAAT promoter element of an MHC class I gene (PD1) results in alternating expression status between different generations

To assess the role of core promoter elements in regulating expression of MHC class I genes, we previously generated a series of transgenic mice with mutations in either the CCAAT box, the TATA box, Sp1 binding site or INR (14) of the swine class I gene, PD1. In that study, we showed that all mutant promoters were capable of supporting expression of the class I transgene. Of 32 independently derived transgenic founder mice with CCAAT element mutations (Fig. 1A), 28 founder mice expressed the transgene at levels equal to or greater than the transgene with a wild type CCAAT box (CCAATwt); the remaining 4 failed to express the PD1 transgene.

**Fig. 1.**
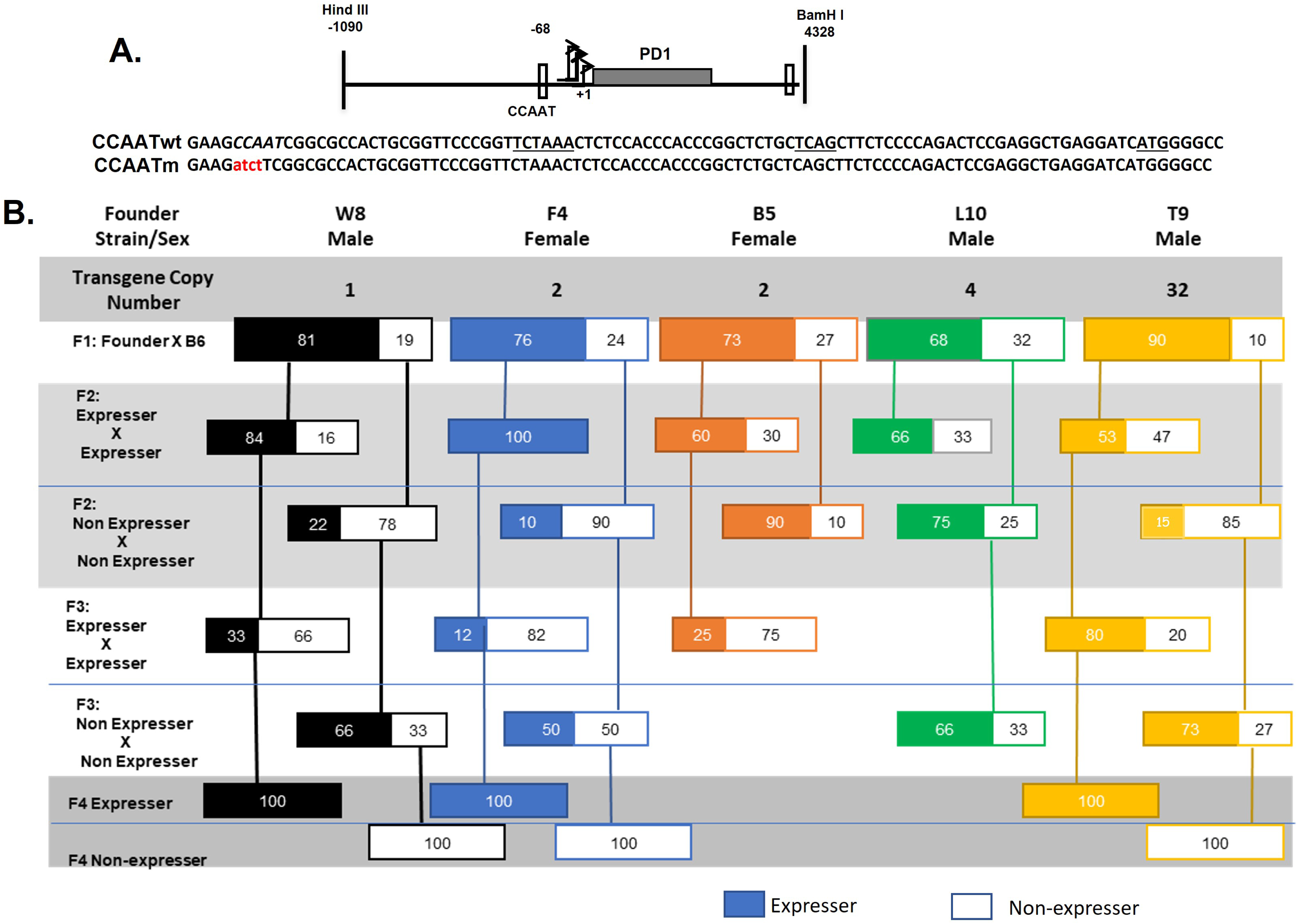
Mutation of the CCAAT box results in variable gene expression across generations. **A.** Both CCAATwt and CCAATm transgenes contain the same backbone DNA segment containing 1kb of 5’ flanking sequences, 3.3kb of MHC class I transgene, PD1, coding sequences (gray rectangle) and 1kb of 3’ sequences. The mutation introduced into the CCAAT sequence (italicized) is indicated; the CCAATwt and mutant CCAAT sequences within the proximal promoter are centered at –68 bp, relative to the major TSS at +1. Transcription of both CCAATwt and CCAAT mutant promoters initiate at the same sites, with a major site at +1. The positions of the TATAA-like element at–30 bp, the Inr at +1, and ATG translation start site are underlined. **B.** Variegated expression of MHC class I transgene, PD1, across multiple generations of transgenic mice with a mutated CCAAT core promoter element. The generational maps are representative of 5 different founder mice with the transgene copy number and sex of the founders indicated. Transgene-positive off-spring were analyzed by FACS for cell surface PD1 expression on PBL. Only transgene-positive mice that express (colored solid boxes) or do not express (outlined, white boxes) are shown.

Five CCAAT mutant founders that expressed the transgene were bred *inter se* to establish lines of CCAAT mutant (CCAATm) transgenes. Pups were monitored for the presence of the PD1 transgene by PCR and for expression of the PD1 protein on peripheral blood lymphocytes (PBL) by flow cytometry. All 5 founders that expressed the transgene gave rise to offspring that also expressed the transgene (Figure 1B). Of note, all 5 lines also generated offspring that carried the transgene but did not express PD1 on their PBL. In subsequent generations of *inter se* breeding, CCAATm mice expressing the PD1 transgene gave rise to offspring that were genotype positive for the transgene and expressed the gene while others did not (Figure 1). Significantly, genotype-positive non-expressers generated both offspring that expressed the transgene and others that did not. All transgene promoter regions were re-sequenced to verify that both expresser and non-expresser transgenes had mutated CCAAT boxes. Additionally, the expression status did not change over the lifetime of an individual mouse.

Thus, expression of the CCAAT mutant transgene flipped between generations. This pattern of variegated expression continued in each independently derived line for multiple generations after which the phenotype was stable and remained stable in all subsequent generations. The variegated expression is unlikely due to insertion site effects, since it was observed in 5 independently derived founders. It is also unlikely due to copy number effects, since three of the lines had low (1–2) copy numbers (Figure 1).Transgenerational variegation of transgene expression was restricted to the CCAATm mice and was not observed in any of the 48 lines of PD1 mice where the CCAAT box was not mutated (these transgenic lines included mice with CCAATwt PD1 transgene as well as with other core promoter mutants i.e. in the TATA box, SP1 binding site or INR mutants) (14).

All further investigations were done in mice that had fixed their expression status as either expressers or non-expressers i.e., after the 4^th^ generation. As shown previously with the CCAATm founder mice that expressed PD1 (14), the CCAATm mice that stably express PD1 do so in all tissues analyzed. Thus, CCAATm transgenic mice that expressed the PD1 protein on their PBL (Fig 2A) also expressed RNA and protein in their tissues (i.e., spleen, kidney, brain) (Figure 2B,C). And as noted previously (14), expression of PD1 in the kidney and brain of CCAATm mice is significantly higher than in CCAATwt mice indicating that the CCAAT box regulates tissue-specific transcription (Fig. 2B,C). Specifically, the CCAAT box appears to negatively regulate class I transcription in non-lymphoid tissues. CCAATm non-expressers that that did not express PD1 protein on their PBL, did not express either PD1 protein or RNA in these tissues (Fig. 2A–C).

**Fig. 2.**
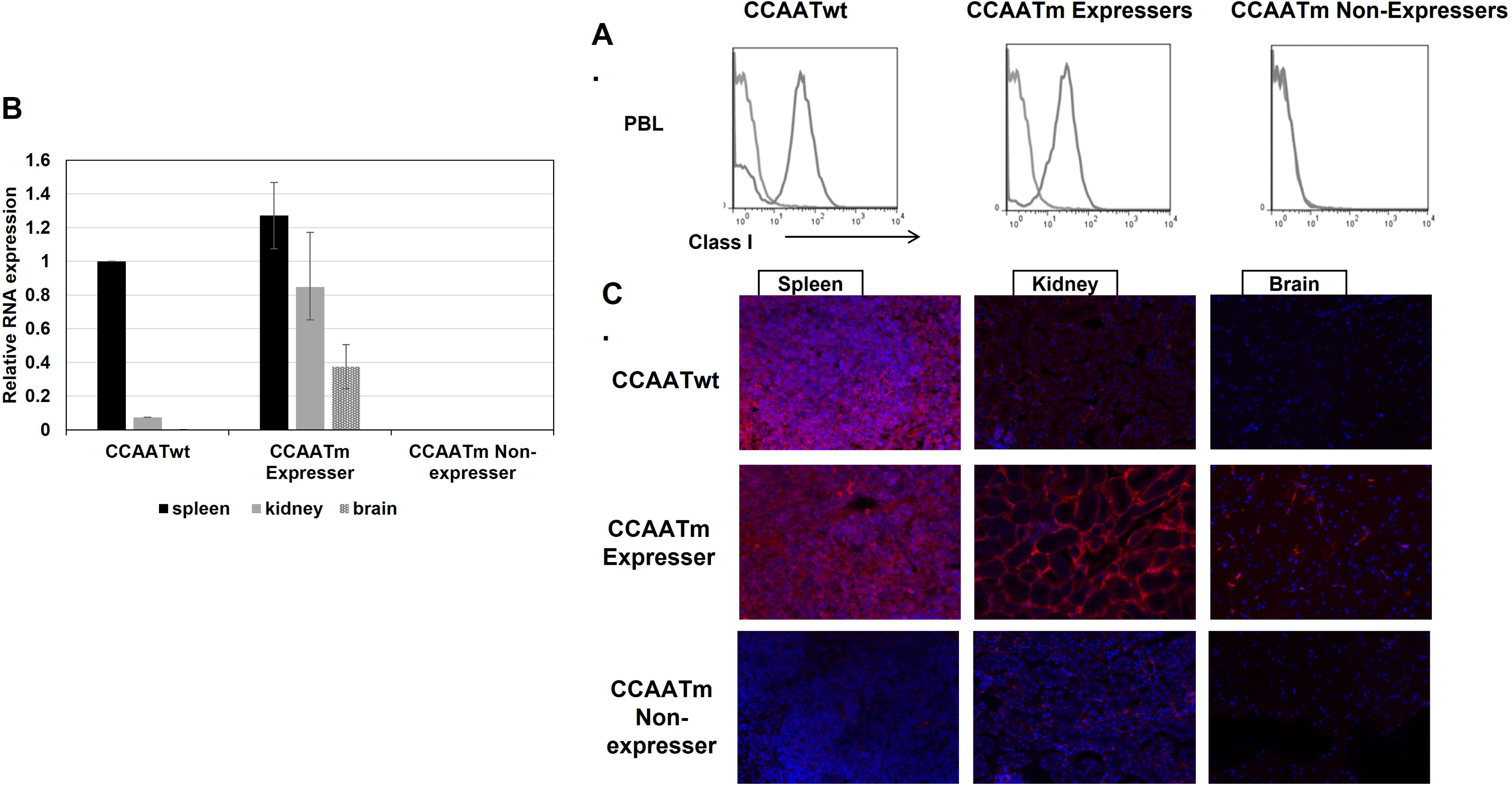
CAATm transgenic mice differ in their MHC class I expression in multiple tissues. **A.** Representative FACS profiles of PBL of the three transgenic lines. PBL were stained with an anti-PD1 antibody (gray profile); secondary antibody alone served as the negative control (light gray profile). Data are representative of >3 independent experiments. **B.** Real-time qPCR was performed on total RNA extracted from tissues of CCAATwt and both CCAAT mutant transgenic lines. The levels of RNA were normalized to 18S RNA in each tissue. Data are average±SEM from 3 individuals of each line. RNA levels in the tissues were all normalized to the RNA level in spleen of CCAATwt mice which was set to 1. **C.** Frozen sections of spleen, kidney and brain from CCAATwt, CCAATm expresser and CCAATm non-expresser transgenic mice were immunostained with anti-PD1 antibody and fluorescent goat anti-mouse Ig. Slides were counterstained with DAPI. Images are representative of two independent experiments.

We next considered the possibility that the CCAATm transgenic non-expressers could be induced to express PD1 in response to external stimuli, such as interferon. As expected, *in vivo* γ-interferon treatment increased the levels of PD1 RNA in the tissues of both CCAATwt and CCAATm expressers. Importantly, it did not induce de novo expression of PD1 in the non-expresser CCAATm transgenic mice, although it did enhance endogenous H2-K^b^ expression in these mice (Fig. 1S). Furthermore, mating of non-expresser CCAATm mice, either male or female, to C57/BL mice did not induce expression in the progeny. Therefore, the lack of PD1 expression by the CCAATm non-expresser mice is an intrinsic defect caused by the CCAAT mutation.

Taken together, these findings demonstrate that mutation of the CCAAT box leads to variegated transgenerational expression of the PD1 gene across generations, where expression status of offspring may vary from that of the parents. Over time, the expression status stabilizes and thereafter the CCAATm offspring are stable expressers or non-expressers. To confirm the role of the CCAAT element in maintaining stable transgenerational expression, a second, independent set of PD1 transgenic mice was established with the same CCAAT mutation. These CCAATm transgenic mice displayed the same transgenerational flipping of expression where expressers gave rise to both expressing and non-expressing offspring and vice-versa (Supplementary Figure 2S).

From these results we conclude that the PD1 CCAAT element has a dual function-as a regulator of transcription and as a determinant of stable transgenerational epigenetic inheritance.

### NF-Y regulates MHC class I expression through the CCAAT box

CCAAT box elements are known to be bound by the trimeric NF-Y complex consisting of NF-YA, NF-YB and NF-YC (23,24). The PD1 CCAAT box, centered at –68bp from the transcription start site (TSS), matches the NF-Y binding consensus sequence. To determine whether NF-Y binds to the PD1 CCAAT box, we first performed an Electrophoretic Mobility Shift Assay (EMSA) using a 50 bp fragment of the PD1 promoter spanning the CCAAT box (Fig 3A). Incubation of the PD1 promoter fragment with HeLa nuclear extract resulted in an electrophoretic mobility shift, indicating that proteins in the extract bound the probe. The addition of anti-NF-YA antibody resulted in a supershift, indicating that the complex contains NF-YA. (NF-YB antibody reduced the intensity of the shifted band, suggesting competition for binding the NF-Y.) Importantly, 1000X excess of unlabeled (cold) CCAATwt probe, but not a CCAATm probe, competed away the complex. These results demonstrate the specific binding of NF-Y to the PD1 WT CCAAT box *in vitro*. This was further demonstrated in vivo using crosslinked ChIPs in splenic cells with NF-YB antibody confirming that NF-Y binds to the CCAATwt but not to CCAATm transgene (Fig 3B).

**Fig. 3.**
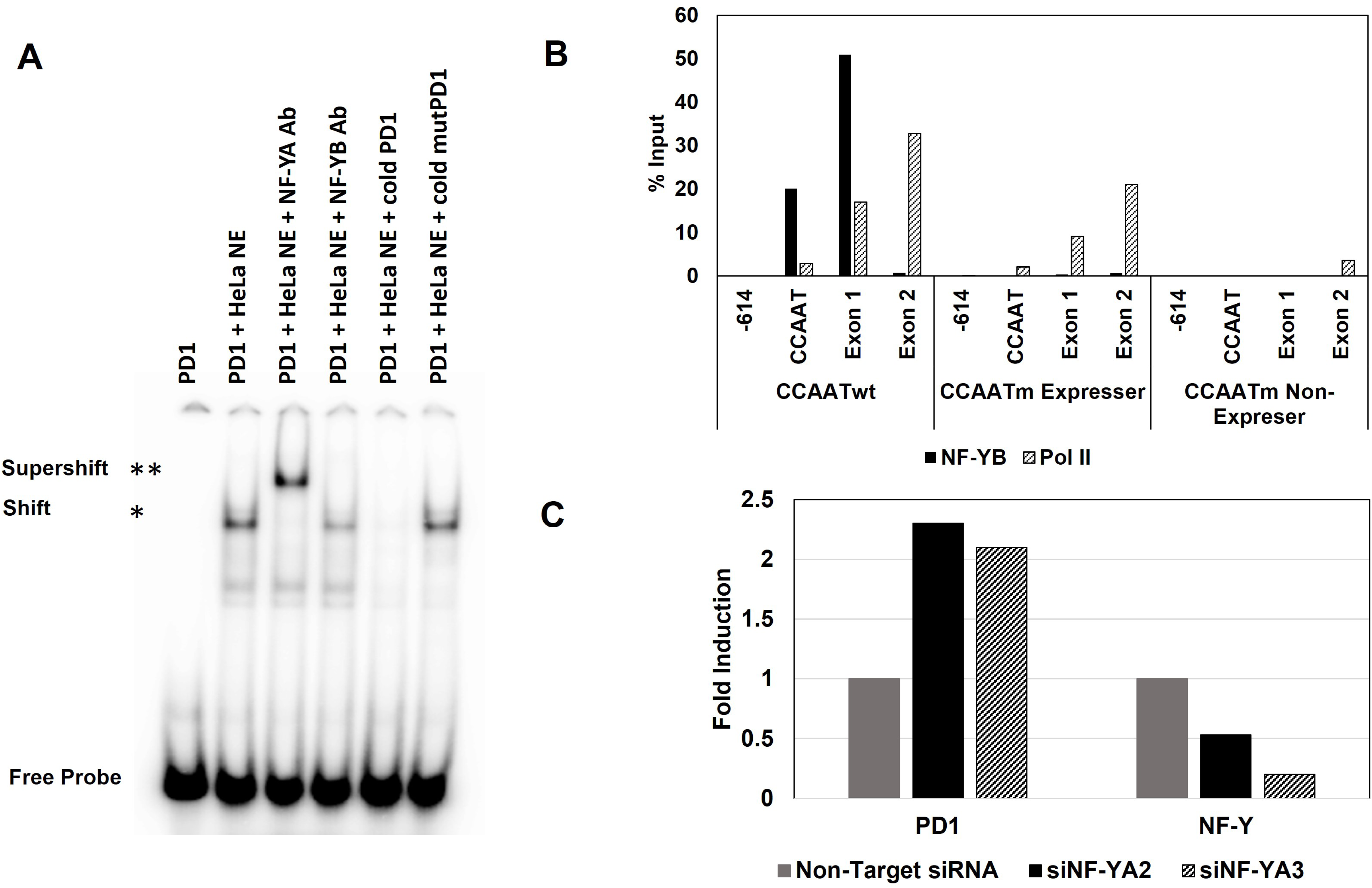
NF-Y binds PD1 CCAAT box both *in vitro* and *in vivo,* and regulates MHC class I expression. **A.** Gel shifts using a radiolabeled PD1 CCAATwt or CCAATm DNA fragment (1011-1060 bp) with or without HeLa nuclear extracts, in the presence or absence of NF-Y antibodies or 1000-fold excess of unlabeled double-stranded CCAATwt or CCAATm oligonucleotide competitors as indicated. **B.** ChIP analysis of NF-YB binding to chromatin from spleens of CCAATwt, CAATm expresser and CAATm non-expresser transgenic strains. Results are expressed as % of total Input. Note: X axis demarcates location relative to the TSS and is not to scale. Data are representative of 2 independent experiments. Pol II ChIP served as positive control where binding was seen in CCAATwt as well as CCAATm expressers but not in CCAATm non-expressers **C.** Quantitation of fold-induction of PD1 gene expression on siRNA knockdown of NF-YA in mouse L cells stably transfected with PD1 (93B2 cell line;NF-YA2 and NF-YA3 were 2 siRNAs targeting NF-YA). The bars labeled PD1 show PD1 RNA levels in cells treated with NF-Y siRNA, expressed relative to the PD1 levels in non-targeting siRNA treated cells set to 1. The bars labeled NF-Y show knock-down of NF-Y RNA levels in NF-Y siRNA treated cells versus cells treated with non-targeting siRNA set to 1. For both PD1 and NF-Y RNA, 18S RNA was used as internal control.

CCAATm expresser mice express higher levels of PD1 RNA and protein in non-lymphoid tissues like kidney and brain as compared to CCAATwt transgenic mice (Fig. 2A and C) (14). The finding that NF-Y binds *in vitro* and *in vivo* to the CCAATwt but not to the mutant CCAAT box, raised the possibility that the increased expression resulted from the absence of NF-Y binding to the mutant CCAAT box *in vivo*. To test this hypothesis, NF-Y was knocked-down in a mouse fibroblast L-cell line stably transfected with the CCAATwt PD1 gene using two siRNAs targeting NF-YA. With either siRNA, depleting NF-YA resulted in increased levels of PD1 transcripts, relative to the non-targeting siRNA control (Figure 3C). These findings are consistent with the aberrantly high expression of the CCAATm transgene in the non-lymphoid cells of CCAATm expressers and demonstrate that NF-Y functions as a negative transcriptional regulator of MHC class I expression in non-lymphoid cells.

### Pol II occupancy correlates with CCAATm transgene expression status

To begin to investigate the basis of transgenerational heterogeneity of expression of PD1 in the CCAAT mutant lines, we first examined Pol II occupancy across the PD1 transgene in splenocytes of CCAATm expressers and non-expressers, as well as CCAATwt transgenic mice. ChIP analysis showed Pol II occupancy at the promoter and across the transgene in both CCAATwt and CCAATm expresser mice (Fig. 4). Overall, the levels of Pol II occupancy across the transgene were comparable between CCAATwt and the CCAATm expressers, except around –150 bp where Pol II levels were markedly higher in CCAATm expressers as compared to CCAATwt mice. In contrast, no Pol II was detected across the body of the PD1 gene of the non-expresser transgene, consistent with the lack of transcription. Surprisingly, Pol II was detected at –75bp of the non-expresser transgene suggesting that although Pol II can load, it cannot progress (Fig. 4).

**Fig. 4.**
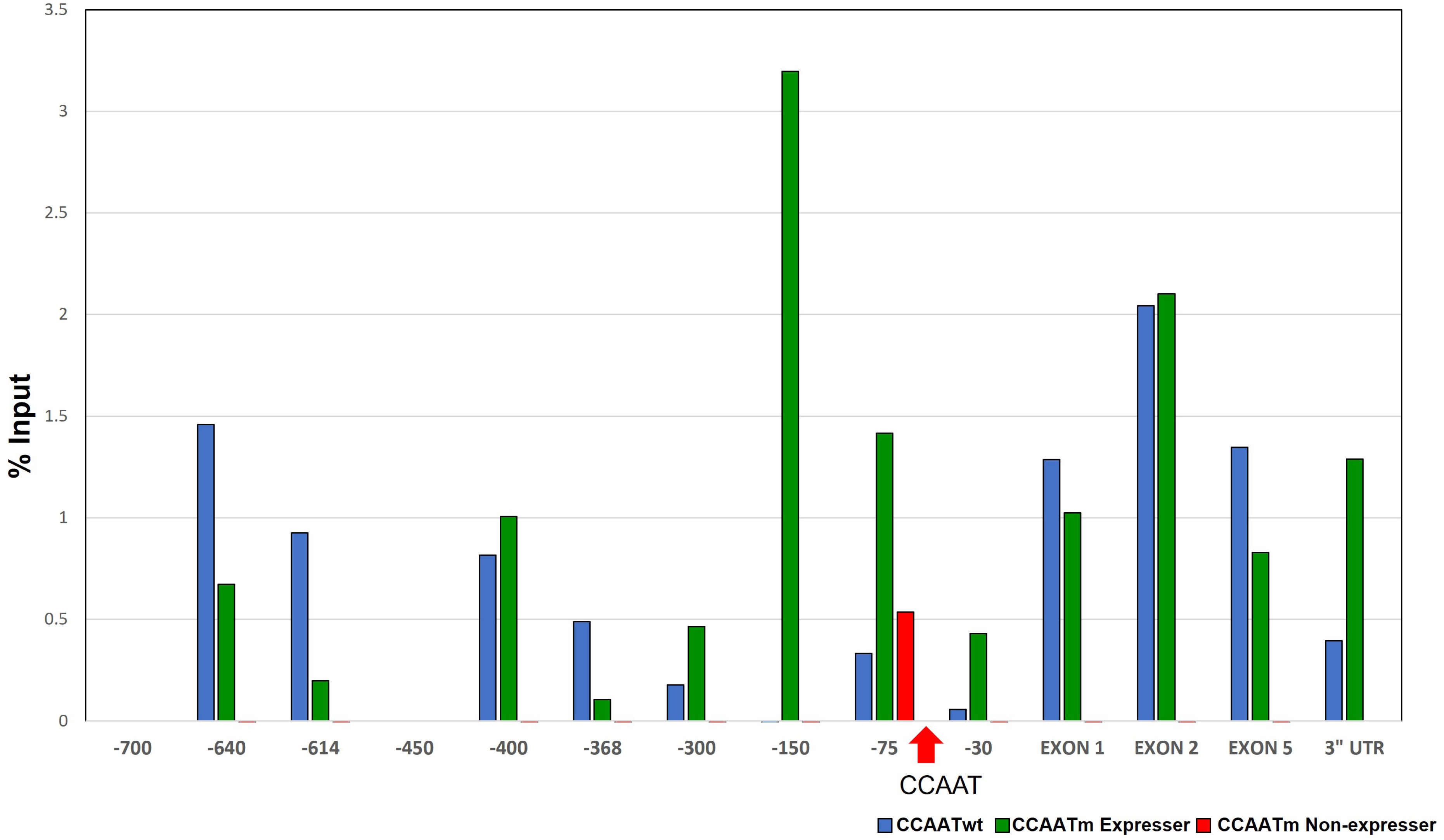
Pol II differs in its association with the CCAATm expresser and non-expresser transgenes. ChIP analysis of Pol II binding to chromatin from spleens of CCAATwt, CAATm expresser and CAATm non-expresser transgenic strains. Results are expressed as % of total Input. Note: X axis demarcates location relative to the TSS and is not to scale. Location of CCAAT box is denoted by arrow. Data are representative of 3 independent experiments.

Pol II occupancy was also detected at distal upstream regulatory regions of the CCAATm expresser, but not the non-expresser, consistent with our observations that the upstream regulatory regions of the PD1 gene are transcribed in PD1 expressing mice but not in the non-expressers (Fig 3S).

Thus, Pol II occupancy at the PD1 gene reflects the expression status of the gene.

### Histone modifications correlate with expression status of the PD1 gene

We next asked whether the differences in Pol II occupancy and expression status of the CCAATm transgenic mice result from differences in chromatin organization or structure. As histone modifications are correlated with epigenetic inheritance, we analyzed the levels of chromatin marks that influence chromatin structure and gene expression. We assessed both histone marks associated with open chromatin and actively transcribing genes (e.g. histone H3 acetylation-H3Ac and histone H3 lysine 4 trimethylation-H3K4me3) and repressive histone chromatin marks (e.g. histone H3 lysine 9 trimethylation-H3K9me3 and H3 lysine 27 trimethylation-H3K27me3), as well as DNA methylation, in spleen cells of the CCAATm and CCAATwt transgenic mice (Fig. 5, 4S and 5S).

**Fig. 5.**
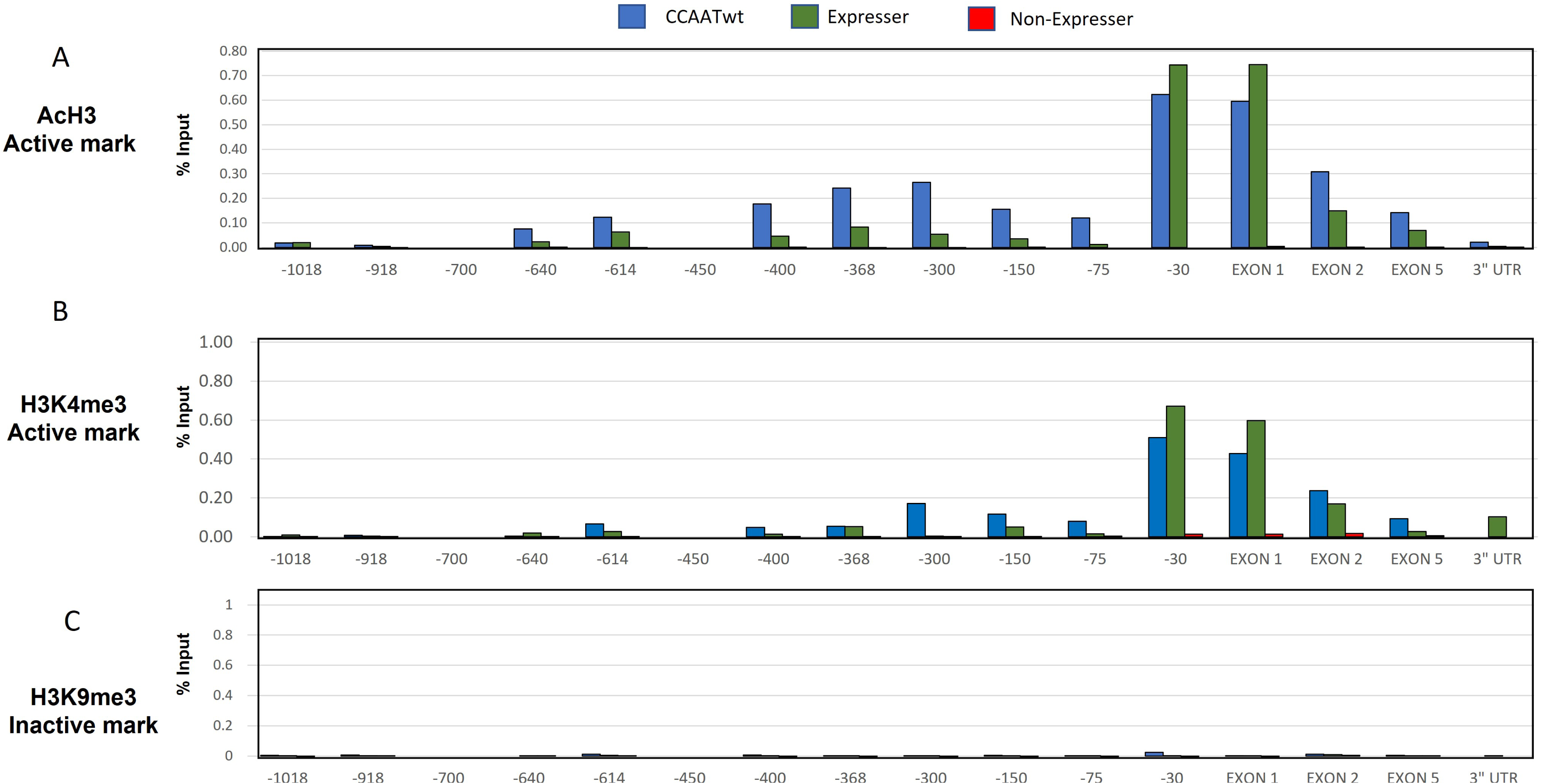
Histone marks of active genes occur only on CCAATwt and CCAATm expressers transgenes. ChIP analysis of AcH3(**A**), H3K4me3 (**B**) and H3K9me3 (**C**) on chromatin from spleens of CCAATwt, expresser, and non-expresser CCAATm transgenic strains. Results are presented as % of total Input. Note: X axis demarcates location relative to the TSS and is not to scale. Data are representative of 3 independent experiments

In splenocytes of CCAATwt and CCAATm expresser mice, the PD1 promoter proximal region and gene body were associated with active histone marks, H3Ac and H3K4me3, across the PD1 transgene (Figure 5A and B). In sharp contrast, the CCAATm non-expresser transgenic mice lacked these positive histone modifications across the PD1 gene. Interestingly, the repressive histone mark H3K9Me3 was not observed across the PD1 transgene in any of these mice, including non-expressers (Figure 5C); H3K27me3 also did not correlate with expression status (Fig. 4SA). Additionally, DNA methylation patterns did not differ between expressers and non-expressers, although both patterns differed from the CCAATwt transgene (Figure 4SB).

In the independently derived second set of CCAATm mice (Fig. 2S), splenic chromatin from CCAATm expresser mice was modified with H3Ac and H3K4me3, but not H3K9me3 (Fig. 5S), as was the original set of CCAATm mice. Chromatin from the CCAATm non-expresser splenocytes was not modified with either H3Ac and H3K4me3, as observed with the original set, but was modified with H3K9me3 marks. The basis for the difference in H3K9me3 pattern between the two sets of mice is not known but may be a reflection of the higher copy number in the second set.

### Nucleosome occupancy does not correlate with PD1 expression status

We reported previously that the CCAATwt PD1 promoter region is nucleosome-free (19), consistent with expression of the transgene. Therefore, we next considered the possibility that the differential expression in CCAATm mice reflected differences in nucleosome occupancy. Nucleosome occupancy across the PD1 transgene was assessed by limiting MNase digestion of nuclei prepared from splenocytes of CCAATwt, CCAATm expressers and non-expressers; DNA recovery was determined by real time PCR. As shown previously, chromatin around the CCAATwt promoter was sensitive to digestion, indicating a relative paucity of nucleosomes across this region (Fig 6). Consistent with the lack of PD1 transcription in the CCAATm non-expresser mice, nucleosome occupancy was much higher across the PD1 promoter. However, unexpectedly, nucleosome occupancy was also high across the PD1 promoter of the CCAATm expresser mice. This finding of similarly high nucleosome occupancy in the expresser transgene was surprising, given its expression status.

**Fig. 6.**
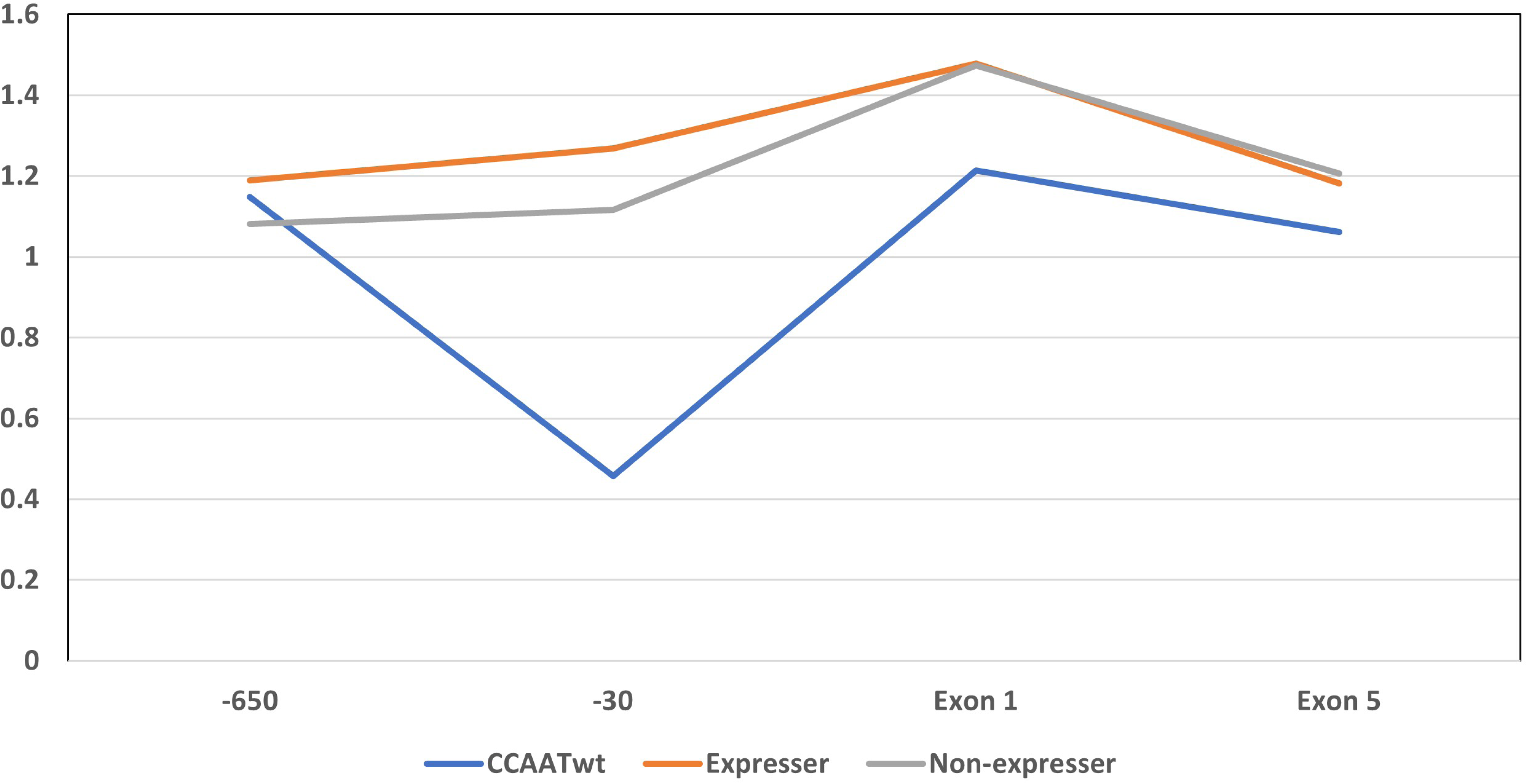
Nucleosome occupancy is higher in both CCAATm transgene lines than in the CCAATwt transgene. Nucleosome occupancy across CCAATm transgene of stable expressers and non-expressers and CCAATwt in spleen. The data include the levels of occupancy at –614, –30, exon 1 and exon 5. The Y axis displays the nucleosomal occupancy following chromatin treatment with 3 units MNase relative to the undigested control. Data are representative of 2 independent experiments.

To examine the possibility that there are subtle differences in nucleosome organization between the CCAATm expressers and non-expressers, chromatin from each was subjected to extensive MNase digestion. The relative susceptibility to digestion of the transgene – a measure of nucleosome occupancy-was assessed by determining DNA recovery in both supernatant and pellet fractions (Fig. 6S). Under conditions of high MNase digestion, the promoters of both CCAATm transgenes were relatively inaccessible compared to the CCAATwt promoter. However, the promoter of the CCAATm expresser transgene was modestly more accessible than the non-expresser, suggesting that this small difference may be sufficient for Pol II loading and elongation.

Thus, although there are small differences in nucleosome occupancy between the CCAATm promoters of expressers and non-expressers, the overall nucleosomal occupancy at the CCAATm promoter does not correlate with expression.

### CCAATm non-expresser mice display a novel pattern of CTCF binding

Although the small difference in chromatin accessibility observed at the promoters of CCAATm expressers and non-expressers might explain in part their patterns of expression, it is likely that some other mechanism establishes and maintains their differential expression. We have previously reported an insulator element associated with the PD1 transgene, located 3’ to the polyA addition site, that is necessary for stable transgene expression (15). We thus considered the possibility that mutation of the CCAAT box altered insulator function associated with the PD1 gene. Since insulator function is often mediated by binding of CTCF (25), we next examined the patterns of CTCF binding to the CCAATwt and CCAAT mutant transgenes in splenic chromatin by ChIP analysis. CTCF binding across the CCAATwt PD1 transgene was centered around –75 bp to –150 bp in the promoter proximal region, just upstream of the CCAAT box (Fig. 7A). CTCF also bound in the same region of the CCAATm expresser proximal to the promoter. No CTCF binding was detected in the distal promoter of either the CCAATwt or CCAATm expresser. In sharp contrast, in the CCAATm non-expresser, a unique CTCF binding site was observed at the 5’ distal promoter, but only low CTCF binding was observed in the promoter proximal region. CTCF also bound within the gene body around exon 5 of both the CCAATm expresser and non-expresser. The promoter distal, but not proximal, binding of CTCF correlates with lack of expression in the CCAATm non-expresser mice. The differential binding of CTCF across the PD1 promoter is independent of cohesin, since cohesin binding was equivalent in the CCAATm lines (Fig. 7S).

**Fig. 7.**
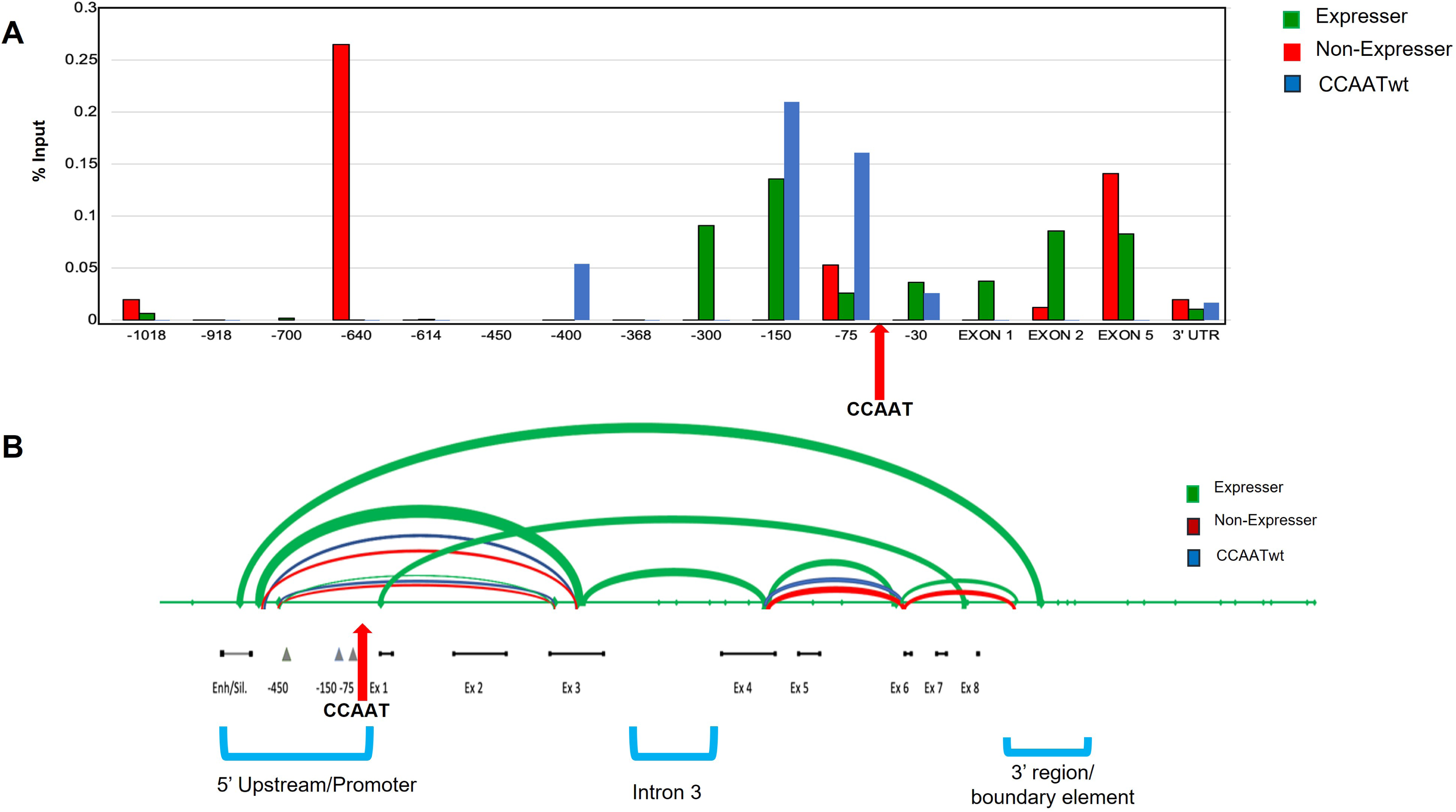
CCAATm transgenes show distinct patterns of CTCF binding and DNA looping. **A.** ChIP analysis of CTCF binding on chromatin from spleens of CCAATwt, expresser and non-expresser CCAATm transgenic strains. Results are presented as % of total Input. Note: X axis denotes location relative to the TSS and is not to scale. Arrow indicates position of the CCAAT box. Data are representative of two independent experiments. **B.** CCAATm expresser transgenes generate DNA loops not seen in either CCAATwt or the CAATm non-expressers transgenes. The endpoints for each loop are located at sites of enzymatic digestion which were ligated as seen by flanking PCRs using the oligos listed in Methods. The thickness of the loop correlates with intensity of the PCR bands. The location of the CCAAT box is denoted by the red arrow. The 5’ Upstream/Promoter, Intron 3, and 3’ region/boundary element are indicated

Thus, the CCAATm non-expressers are distinguished from the expressers and WT by their CTCF binding patterns.

### Distinct DNA looping patterns correlate with PD1 gene expression in CCAATm mice

CTCF is known to mediate both long and short loop formation through dimerization of two CTCF molecules bound at distal sites. Long loop formation is facilitated by cohesin, whereas short loop formation often occurs independently of cohesin and brings together promoter and gene body regions (26,27). The novel upstream binding of CTCF to the non-expresser transgene, in combination with its binding to the 3’ region of the gene, raised the possibility that CTCF forms novel chromatin loops in the non-expresser transgene. Therefore, we examined whether local looping occurs across the PD1 transgene by 3C chromosome capture (Fig. 7B). All 3 transgenes shared two major loops. One loop is centered around the promoter and spans the CCAAT box. Its formation is independent of NF-Y binding since both the WT and CCAATm transgenes form this loop. It also should be noted that the CTCF binding site in the CCAATm non-expresser lies outside of the loop that maps to the 5’ promoter region. Thus, CTCF binding is unlikely to affect formation of this loop. The second common loop is centered between exon 5 and exon 6. CTCF binds to this region in both the CCAATm expresser and non-expresser but not the WT, making it unlikely that it is involved in regulating either loop formation or expression.

Surprisingly, the CCAATm expresser formed an additional loop that is centered around intron 3 and not found in either the CCAATwt or CCAATm non-expresser transgenes (Fig. 7B). What factors contribute to the formation of this loop in the CCAATm expresser remain to be determined. However, these findings suggest that, in the absence of the wild type CCAAT box, the unique loop in the CCAATm expresser gene serves to protect the gene from inactivation.

## Discussion

In the present study, we report that the CCAAT box associated with the MHC class I promoter is a regulator of transgenerational epigenetic inheritance (TEI).

Although TEI has been observed in plants and mammals, the mechanisms underlying TEI are still not known. We have now extended the understanding of TEI by characterizing the role of the MHC class I CCAAT box promoter element in establishing stable expression across generations. Mutation of the CCAAT box results in the failure of NF-Y binding, aberrant patterns of MHC class I expression and the transgenerational instability of expression of the MHC class I transgene, stabilizing only after multiple generations. Thus, the MHC class I CCAAT box functions as both a regulator of transcription and to maintain stable gene expression across generations.

Among the mechanisms maintaining epigenetic inheritance in plants, the retention of DNA methylation and nucleosomal packaging during embryogenesis have been proposed (1). Although both of these mechanisms have been reported, they play only a minor role in TEI in mammals (28–31). In the MHC class I CCAATm transgene, expression correlates with the presence of H3K4me3 and total H3 acetylation, but it does not correlate with DNA methylation. NF-Y binding previously has been shown to affect post-translation modifications on chromatin histones through its interaction with different chromatin modifiers or remodeling complexes (32). However, MHC class I expression status in the CCAATm transgenes was independent of the ability to bind NF-Y. Similarly, we observed only minor differences in nucleosomal packaging between expressers and non-expressers. Thus, these two potential mechanisms are not operational for TEI associated with the CCAAT box mutant.

Given the large-scale epigenetic reprograming in mammals during embryogenesis, other mechanisms – such as non-coding RNAs, 3D genome organization, and transcription factor binding-may contribute to mammalian TEI (1). Indeed, all three of these features distinguish the CCAATm expresser from the non-expressers. Whereas the CCAATm expresser transgene transcribes promoter-distal non-coding RNA sequences, the CCAATm non-expresser does not. However, whether this is the mechanism determining TEI or a reflection of the ability of Pol II to initiate transcription cannot be distinguished. Significantly, the CCAATm expressers and non-expressers differ both in their 3D genome organization and transcription factor binding. The CCAATm expresser forms a unique 3D loop not found in either the CCAATm non-expresser or the WT transgene. This loop encompasses exon 3, intron 3 and exon 4 which does not span any of the regions bound by CTCF. The role of this loop in expression of the CCAATm transgene, as well as its precise anchors, remain to be determined. Conversely, CTCF binds to a region in the 5’ distal promoter of the CCAATm non-expresser, but not to the expresser. We speculate that CTCF binding to this site acts as a barrier to expression, perhaps by loop formation with the previously-identified 3’ barrier element (15), although we have not been able to detect such a loop to date. Thus, there is a reciprocal relationship between CTCF binding and loop formation that correlates with expression; whether the establishment of one prevents the other remains to be determined. These findings extended previous reports that allele-specific differences in CTCF binding can lead to altered chromatin environment, 3D structure and gene expression (33). The variegated patterns of PD1 expression that occur in the early CCAATm transgene generations reflect stochastic establishment of either loops or CTCF binding. What leads to a stable, heritable configuration after the early generations remains to be determined, as does the role of DNA sequence elements.

The present studies suggest that the CCAAT box plays a singular role in the stable expression of the MHC class I gene. Thus, none of the forty-eight transgenic lines generated from either the WT MHC class I gene or other MHC class I promoter mutants (mutants in the TATA box, SP1 BS or Inr) display the variegated expression observed in the multiple CCAATm transgenic lines. There is only limited previous evidence for the role of a DNA sequence element in regulating TEI. DNA methylation of CpG islands associated with promoters is prevented by the presence of a promoter-associated Sp1-binding site; whether Sp1 is the responsible transcription factor was not directly demonstrated (34). In the present study, mutation of the CCAAT box abrogates NF-Y binding, further suggesting that NF-Y plays a critical role in maintaining stable expression. We speculate that the mechanism leading to the initiation of TEI upon mutation of the CCAATm box results from loss of NF-Y binding leading to aberrant looping and transcription factor binding. In wild-type animals, NF-Y binding presumably establishes a favorable 3D chromatin structure that results in stable expression of the Class I gene across generations. NF-Y has been shown to contribute to zygotic genome activation and formation of DNase hypersensitive sites at the 2-cell stage (35). So, loss of NF-Y binding in CCAAT mutants presumably results in stochastic expression very early in embryogenesis leading to differential expression in littermates. This expression status is maintained throughout the lifespan of the individual but is reprogrammed and reset in subsequent generations.

As Rothi and Greer (7) pointed out in a recent review, the understanding of TEI now needs to go beyond correlation to causation, about which relatively little is known. Recent studies in C. elegans have begun to track heritable molecules that transmit epigenetic information. In worms, viral infections and starvation result in transgenerational inheritance of small RNAs that influence the transcriptome (36,37). Unlike previous examples of transgenerational inheritance which result from environmental triggers, these studies define a novel mechanism of epigenetic inheritance caused by a discrete mutation within a transcription factor binding site, leading to formation of novel DNA loops. Our demonstration, in multiple independent transgenic lines, that mutation of the CCAAT box results in TEI provides a clear example of causation in mammals.

## ACKNOWLEDGEMENTS

The authors gratefully acknowledge other members of the Singer lab for helpful discussions, the lab of Yamini Dalal, Stanley Adoro and Christian Mayer for their critical reading of the manuscript. This research was supported by the Intramural Research Program of the NIH, National Cancer Institute, Center for Cancer Research.

## DATA AVAILABILITY

All reagents, resources and data reported in this manuscript are freely available.

## REFERENCES

1. Fitz-James, M.H. and Cavalli, G. (2022) Molecular mechanisms of transgenerational epigenetic inheritance. Nat Rev Genet, 23, 325–341.

2. Cavalli, G. and Heard, E. (2019) Advances in epigenetics link genetics to the environment and disease. Nature, 571, 489–499.

3. Arzate-Mejia, R.G. and Mansuy, I.M. (2022) Epigenetic Inheritance: Impact for Biology and Society-recent progress, current questions and future challenges. Environ Epigenet, 8, dvac021.

4. Burton, N.O. and Greer, E.L. (2022) Multigenerational epigenetic inheritance: Transmitting information across generations. Semin Cell Dev Biol, 127, 121–132.

5. Heard, E. and Martienssen, R.A. (2014) Transgenerational epigenetic inheritance: myths and mechanisms. Cell, 157, 95–109.

6. Inoue, A., Jiang, L., Lu, F., Suzuki, T. and Zhang, Y. (2017) Maternal H3K27me3 controls DNA methylation-independent imprinting. Nature, 547, 419–424.

7. Rothi, M.H. and Greer, E.L. (2023) From correlation to causation: The new frontier of transgenerational epigenetic inheritance. Bioessays, 45, e2200118.

8. Howcroft, T. and Singer, D. (2003) Expression of nonclassical MHC class Ib genes: comparison of regulatory elements. Immunol Res, 27, 1–30.

9. van den Elsen, P.J. (2011) Expression regulation of major histocompatibility complex class I and class II encoding genes. Front Immunol, 2, 48.

10. Louis-Plence, P., Moreno, C.S. and Boss, J.M. (1997) Formation of a regulatory factor X/X2 box-binding protein/nuclear factor-Y multiprotein complex on the conserved regulatory regions of HLA class II genes. J Immunol, 159, 3899–3909.

11. Martyn, G.E., Quinlan, K.G.R. and Crossley, M. (2017) The regulation of human globin promoters by CCAAT box elements and the recruitment of NF-Y. Biochim Biophys Acta Gene Regul Mech, 1860, 525–536.

12. Mantovani, R., Pessara, U., Tronche, F., Li, X.Y., Knapp, A.M., Pasquali, J.L., Benoist, C. and Mathis, D. (1992) Monoclonal antibodies to NF-Y define its function in MHC class II and albumin gene transcription. EMBO J, 11, 3315–3322.

13. Frels, W.I., Bluestone, J.A., Hodes, R.J., Capecchi, M.R. and Singer, D.S. (1985) Expression of a microinjected porcine class I major histocompatibility complex gene in transgenic mice. Science, 228, 577–580.

14. Barbash, Z.S., Weissman, J.D., Campbell, J.A., Jr., Mu, J. and Singer, D.S. (2013) Major histocompatibility complex class I core promoter elements are not essential for transcription in vivo. Mol Cell Biol, 33, 4395–4407.

15. Cohen, H., Parekh, P., Sercan, Z., Kotekar, A., Weissman, J.D. and Singer, D.S. (2009) In vivo expression of MHC class I genes depends on the presence of a downstream barrier element. PLoS One, 4, e6748.

16. Ehrlich, R., Sharrow, S.O., Maguire, J.E. and Singer, D.S. (1989) Expression of a class I MHC transgene: effects of in vivo alpha/beta-interferon treatment. Immunogenetics, 30, 18–26.

17. Weissman, J.D. and Singer, D.S. (1991) A complex regulatory DNA element associated with a major histocompatibility complex class I gene consists of both a silencer and an enhancer. Mol Cell Biol, 11, 4217–4227.

18. Bernardini, A., Lorenzo, M., Nardini, M., Mantovani, R. and Gnesutta, N. (2019) The phosphorylatable Ser320 of NF-YA is involved in DNA binding of the NF-Y trimer. FASEB J, 33, 4790–4801.

19. Kotekar, A.S., Weissman, J.D., Gegonne, A., Cohen, H. and Singer, D.S. (2008) Histone modifications, but not nucleosomal positioning, correlate with major histocompatibility complex class I promoter activity in different tissues in vivo. Mol Cell Biol, 28, 7323–7336.

20. Landel, C.P., Stabley, D.L. and Bundesen, L.Q. (1997) PCR identification of class I major histocompatibility complex genes transcribed in mouse blastocyst and placenta. Journal of reproductive immunology, 33, 31–43.

21. Splinter, E., Grosveld, F. and de Laat, W. (2004) 3C technology: analyzing the spatial organization of genomic loci in vivo. Methods Enzymol, 375, 493–507.

22. Cope, N.F. and Fraser, P. (2009) Chromosome conformation capture. Cold Spring Harb Protoc, 2009, pdb prot5137.

23. Bi, W., Wu, L., Coustry, F., de Crombrugghe, B. and Maity, S.N. (1997) DNA binding specificity of the CCAAT-binding factor CBF/NF-Y. J Biol Chem, 272, 26562–26572.

24. Maity, S.N., Sinha, S., Ruteshouser, E.C. and de Crombrugghe, B. (1992) Three different polypeptides are necessary for DNA binding of the mammalian heteromeric CCAAT binding factor. J Biol Chem, 267, 16574–16580.

25. Cuddapah, S., Jothi, R., Schones, D.E., Roh, T.Y., Cui, K. and Zhao, K. (2009) Global analysis of the insulator binding protein CTCF in chromatin barrier regions reveals demarcation of active and repressive domains. Genome Res, 19, 24–32.

26. Merkenschlager, M. and Odom, D.T. (2013) CTCF and cohesin: linking gene regulatory elements with their targets. Cell, 152, 1285–1297.

27. Ruiz-Velasco, M., Kumar, M., Lai, M.C., Bhat, P., Solis-Pinson, A.B., Reyes, A., Kleinsorg, S., Noh, K.M., Gibson, T.J. and Zaugg, J.B. (2017) CTCF-Mediated Chromatin Loops between Promoter and Gene Body Regulate Alternative Splicing across Individuals. Cell Syst, 5, 628–637 e626.

28. Brykczynska, U., Hisano, M., Erkek, S., Ramos, L., Oakeley, E.J., Roloff, T.C., Beisel, C., Schubeler, D., Stadler, M.B. and Peters, A.H. (2010) Repressive and active histone methylation mark distinct promoters in human and mouse spermatozoa. Nat Struct Mol Biol, 17, 679–687.

29. Gold, H.B., Jung, Y.H. and Corces, V.G. (2018) Not just heads and tails: The complexity of the sperm epigenome. J Biol Chem, 293, 13815–13820.

30. Hammoud, S.S., Nix, D.A., Zhang, H., Purwar, J., Carrell, D.T. and Cairns, B.R. (2009) Distinctive chromatin in human sperm packages genes for embryo development. Nature, 460, 473–478.

31. Morgan, H.D., Santos, F., Green, K., Dean, W. and Reik, W. (2005) Epigenetic reprogramming in mammals. Hum Mol Genet, 14 **Spec No 1**, R47-58.

32. Dolfini, D., Gatta, R. and Mantovani, R. (2012) NF-Y and the transcriptional activation of CCAAT promoters. Crit Rev Biochem Mol Biol, 47, 29–49.

33. McDaniell, R., Lee, B.K., Song, L., Liu, Z., Boyle, A.P., Erdos, M.R., Scott, L.J., Morken, M.A., Kucera, K.S., Battenhouse, A. et al. (2010) Heritable individual-specific and allele-specific chromatin signatures in humans. Science, 328, 235–239.

34. Brandeis, M., Frank, D., Keshet, I., Siegfried, Z., Mendelsohn, M., Nemes, A., Temper, V., Razin, A. and Cedar, H. (1994) Sp1 elements protect a CpG island from de novo methylation. Nature, 371, 435–438.

35. Lu, F., Liu, Y., Inoue, A., Suzuki, T., Zhao, K. and Zhang, Y. (2016) Establishing Chromatin Regulatory Landscape during Mouse Preimplantation Development. Cell, 165, 1375–1388.

36. Rechavi, O., Houri-Ze’evi, L., Anava, S., Goh, W.S.S., Kerk, S.Y., Hannon, G.J. and Hobert, O. (2014) Starvation-induced transgenerational inheritance of small RNAs in C. elegans. Cell, 158, 277–287.

37. Rechavi, O., Minevich, G. and Hobert, O. (2011) Transgenerational inheritance of an acquired small RNA-based antiviral response in C. elegans. Cell, 147, 1248–1256.

