## Supplemental Figures for "Transgenerational Epigenetic Inheritance of MHC Class I Gene Expression is Regulated by the CCAAT Promoter Element"

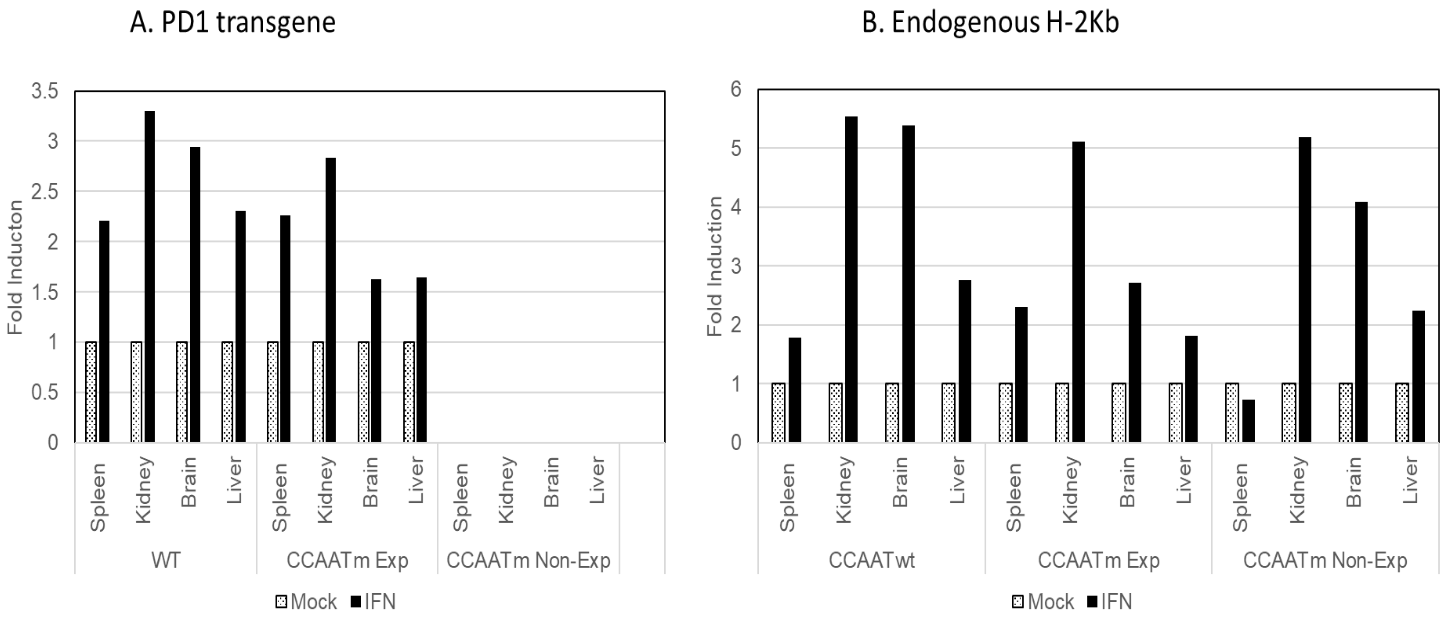


**Fig. 1S: CCAATm Expressers, but Not Non-expressers, Respond to γ-Interferon Treatment**

Mice were mock-treated or treated with γ-Interferon; tissues were harvested 24 hours post treatment. Stipple boxes: Mock treated. Black boxes: γ-Interferon treated. A: PD1 RNA levels. B: $H2K^b$ RNA levels. Data are presented relative to the CCAATwt control, normalized to 18S.

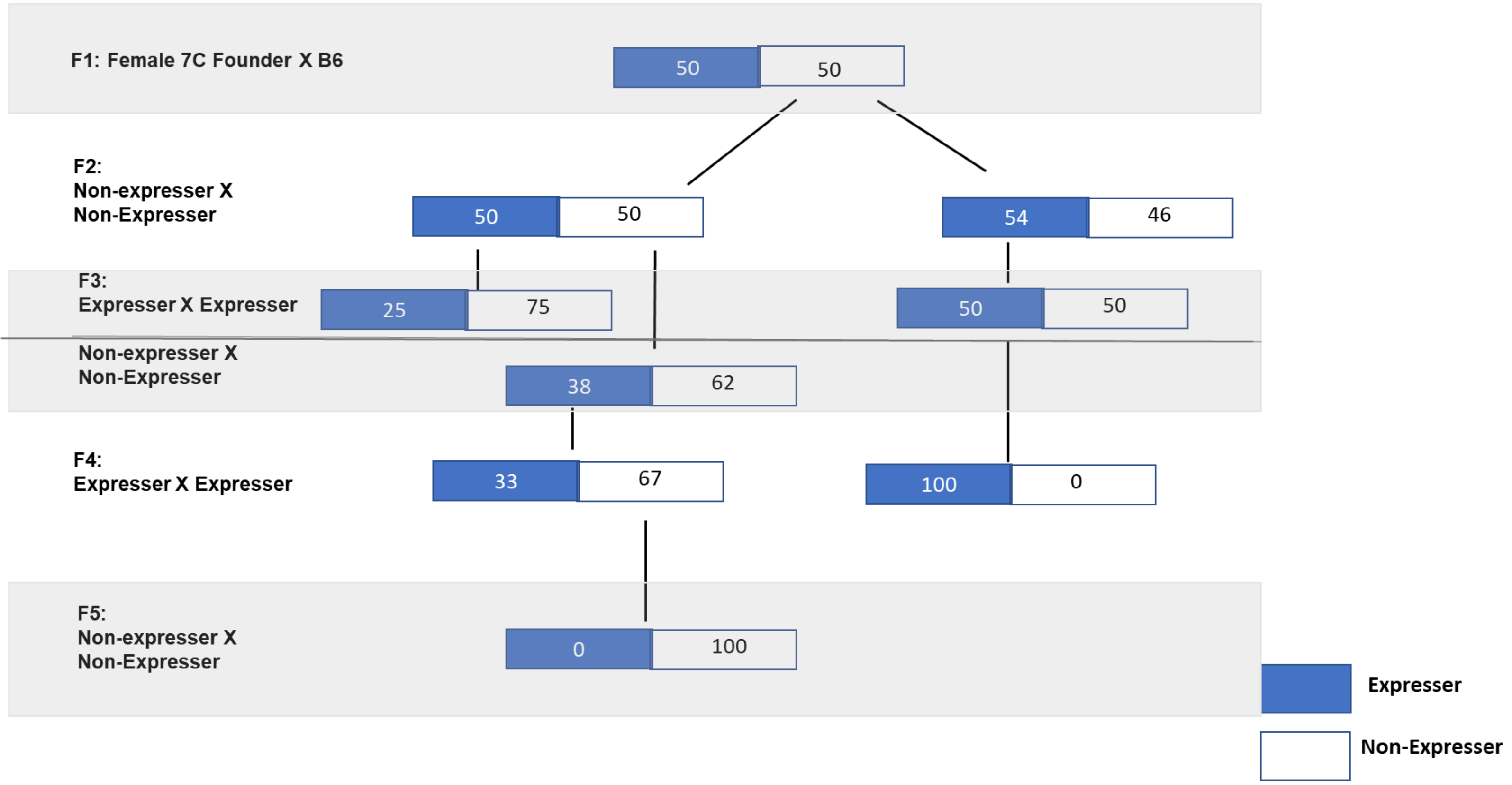


**Fig. 2S: Transgenerational Epigenetic Inheritance in an Independent Series of Transgenic Mice**

Variegated expression of MHC class I, PD1, across multiple generations of an independent transgenic mouse line (7C) with a mutated CCAAT core promoter element and a copy number of 24. Transgene-positive off-spring were analyzed by FACS for cell surface PD1 expression on PBL. Only transgene-positive mice that express (solid boxes) or do not express (outlined, white boxes) are shown.

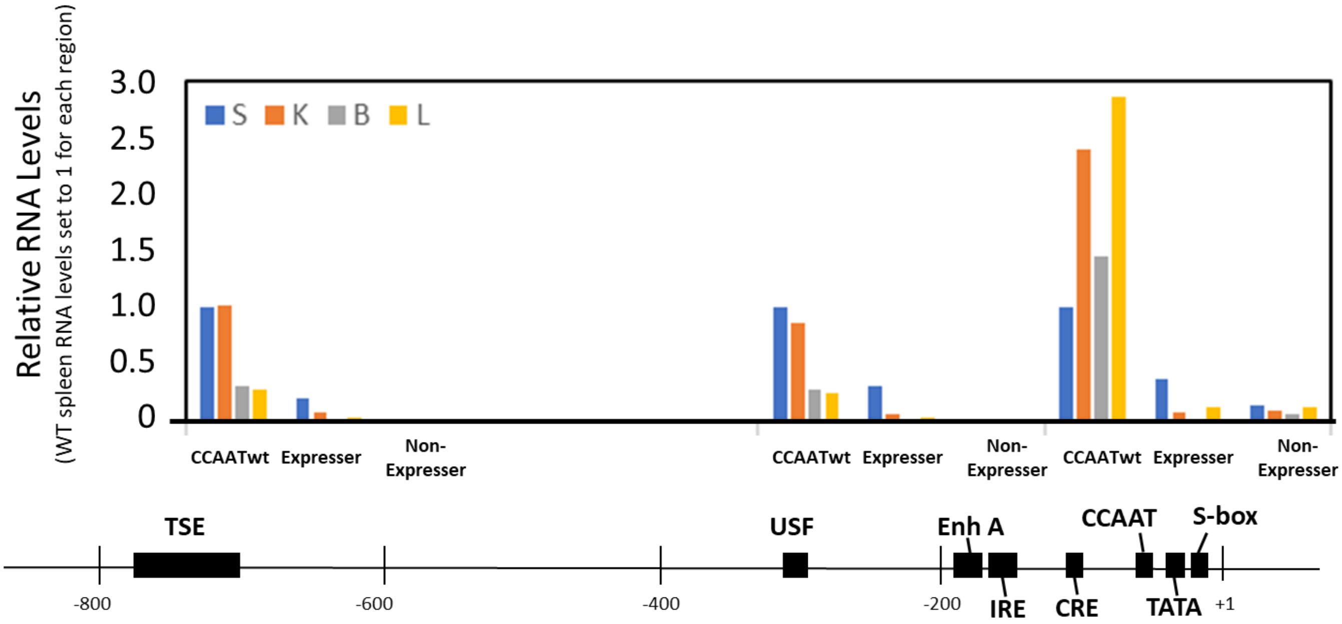


**Fig 3S: Upstream Regulatory Regions of the PD1 Gene are Transcribed in CCAAT Mutant Expressers, but not Non-Expressers**

RNA levels at upstream start sites are shown in tissues from CCAAT mutant expresser and non-expresser mouse strains. CCAATwt spleen RNA levels are set to 1 for each region. Location of upstream PCR primers relative to +1 start site are indicated under the X-Axis as well as in the map below.

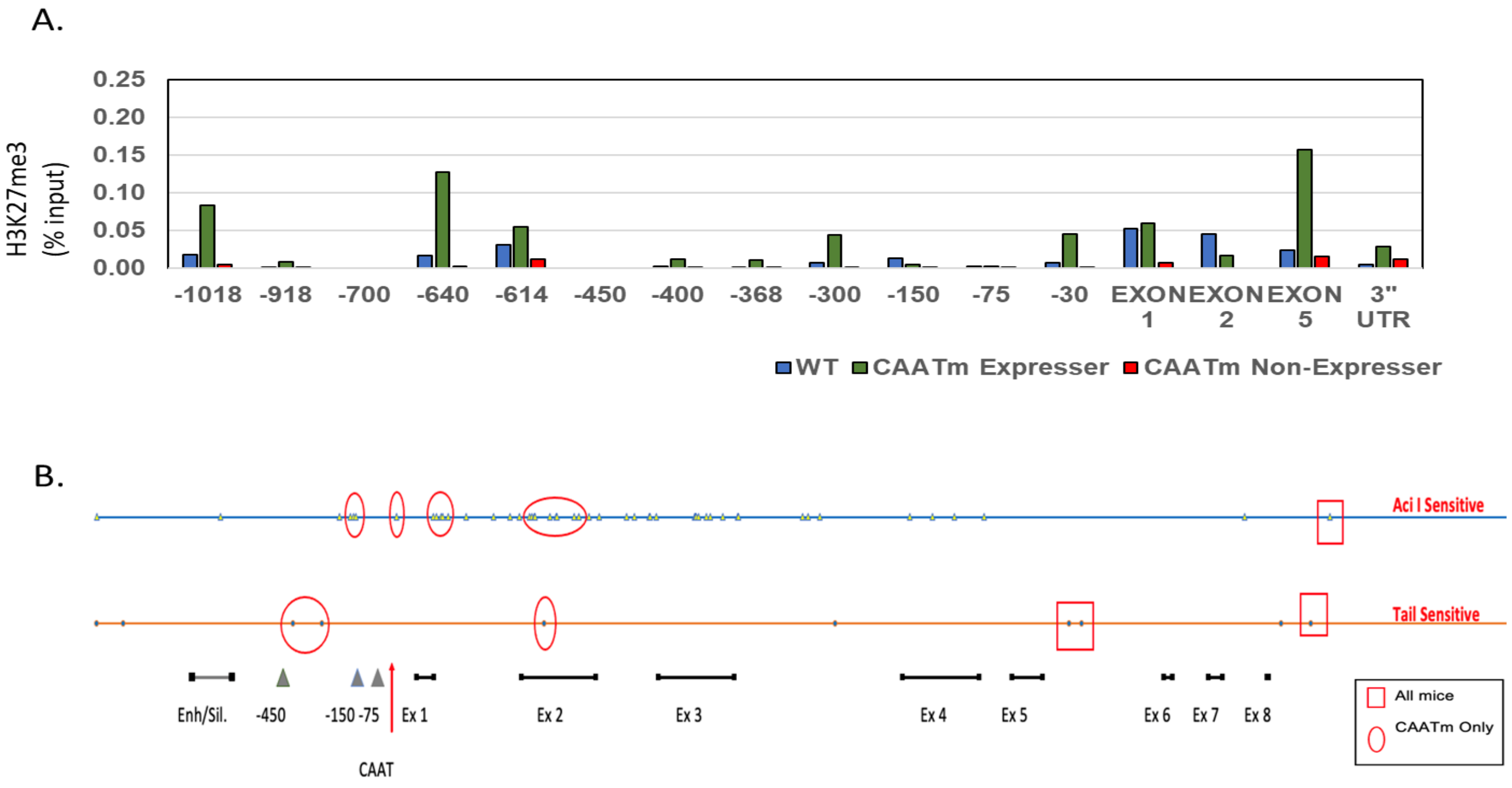


**Fig. 4S: DNA and H3K27 Methylation Patterns Do Not Distinguish CCAATm Expressers from Non-Expressers**

**A.** ChIP analysis of H3K27me3 binding to chromatin across transgene from spleens of CCAATwt, CCAATm expresser and CCAATm non-expresser mice as % of total Input. Note X axis denotes location relative to the TSS and is not to scale.

**B.** DNA methylation analysis across transgene of CCAATwt, CCAATm expresser and CCAATm non-expresser mice. Restriction enzyme sites for methylation sensitive enzymes Aci 1 (dots, upper line) and Tai1 (triangles, lower line) are indicated. Open oval symbols represent locations of DNA methylation in the CCAAT mutant strains. Open boxes represent locations of DNA methylation in all strains. Below is schematic location of upstream regulatory elements and Exons relative to the enzyme sites.

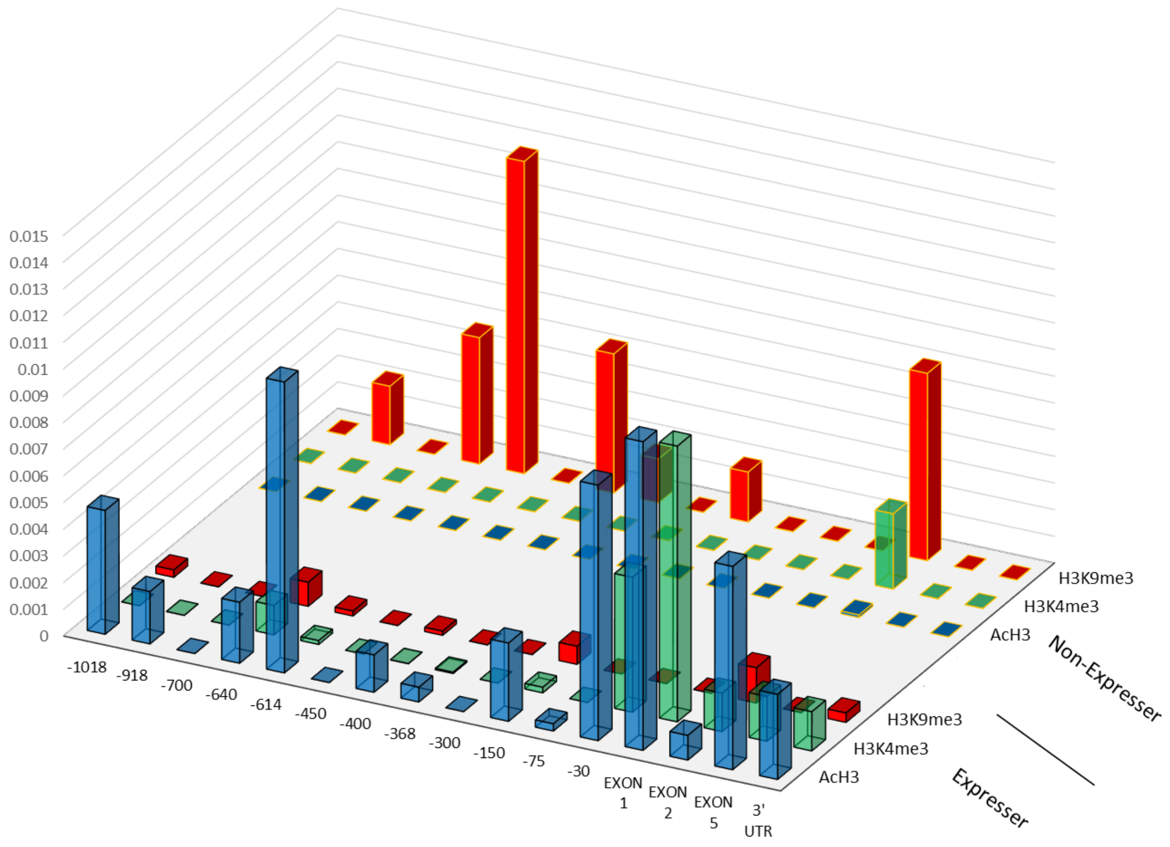


**Fig 5S: Histone marks associated with CCAATm transgenes in lines derived from a single independent CCAATm founder correlates with expression.**

ChIP analysis of AcH3, H3K4me3, H3K9me binding to chromatin from spleens of CCAATm expresser and CCAATm non-expresser transgenics (line 7C) which share a common founder, Results are expressed as % of total Input. Note: X axis denotes location relative to the TSS and is not to scale.

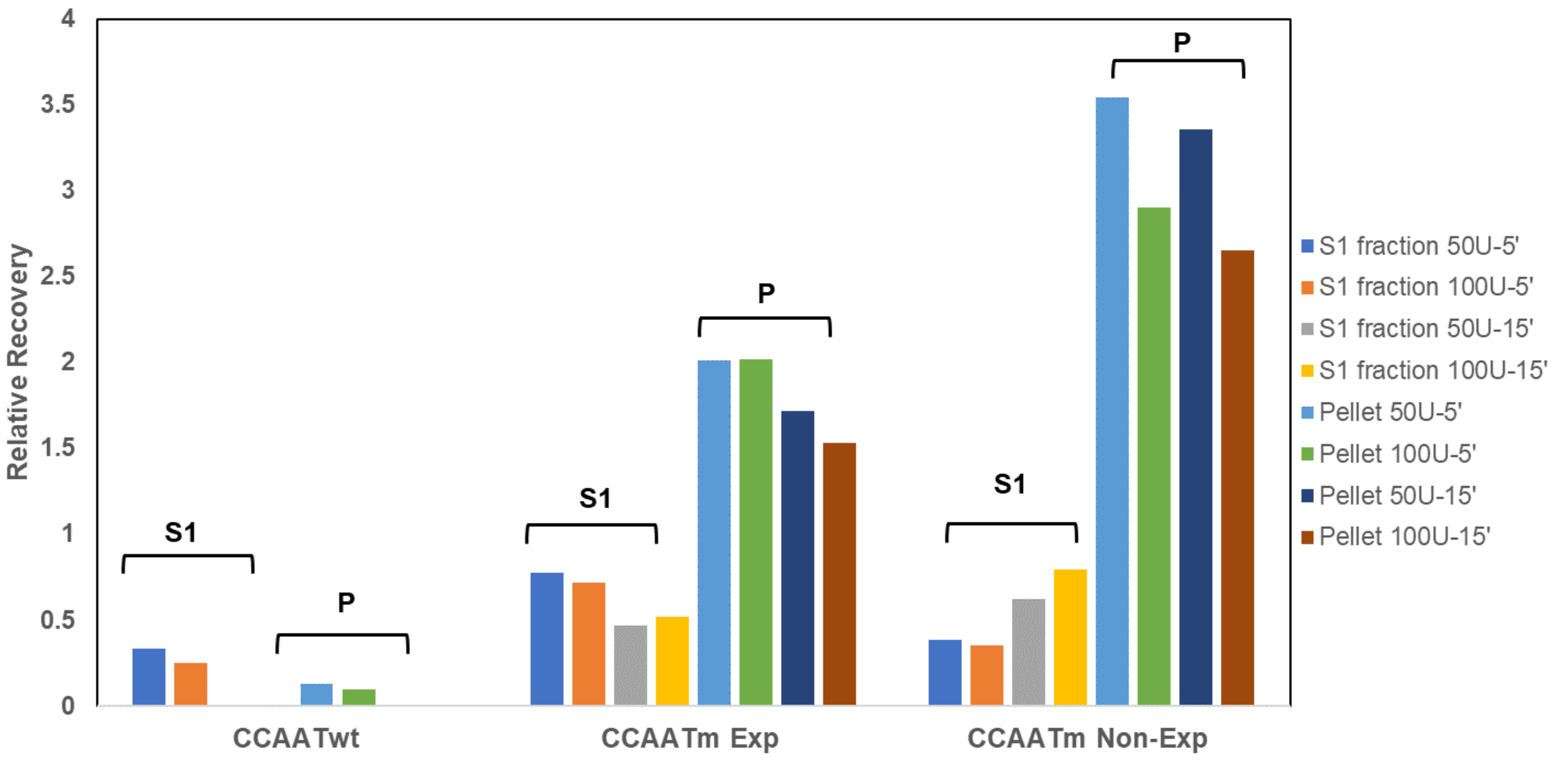


**Figure 6S: PD1 Promoter DNA is Relatively Inaccessible in CCAAT Mutants Relative to CCAATwt**

Nuclei from spleens of CCAATwt, CCAATm expresser and CCAATm non-expresser mice were digested with 50 or 100 units of MNase and recovery of DNA in the supernatants and pellets were accessed for quantity and location. S, K, B and L represent data from spleen, kidney, brain and liver, respectively. Results are from one experiment.

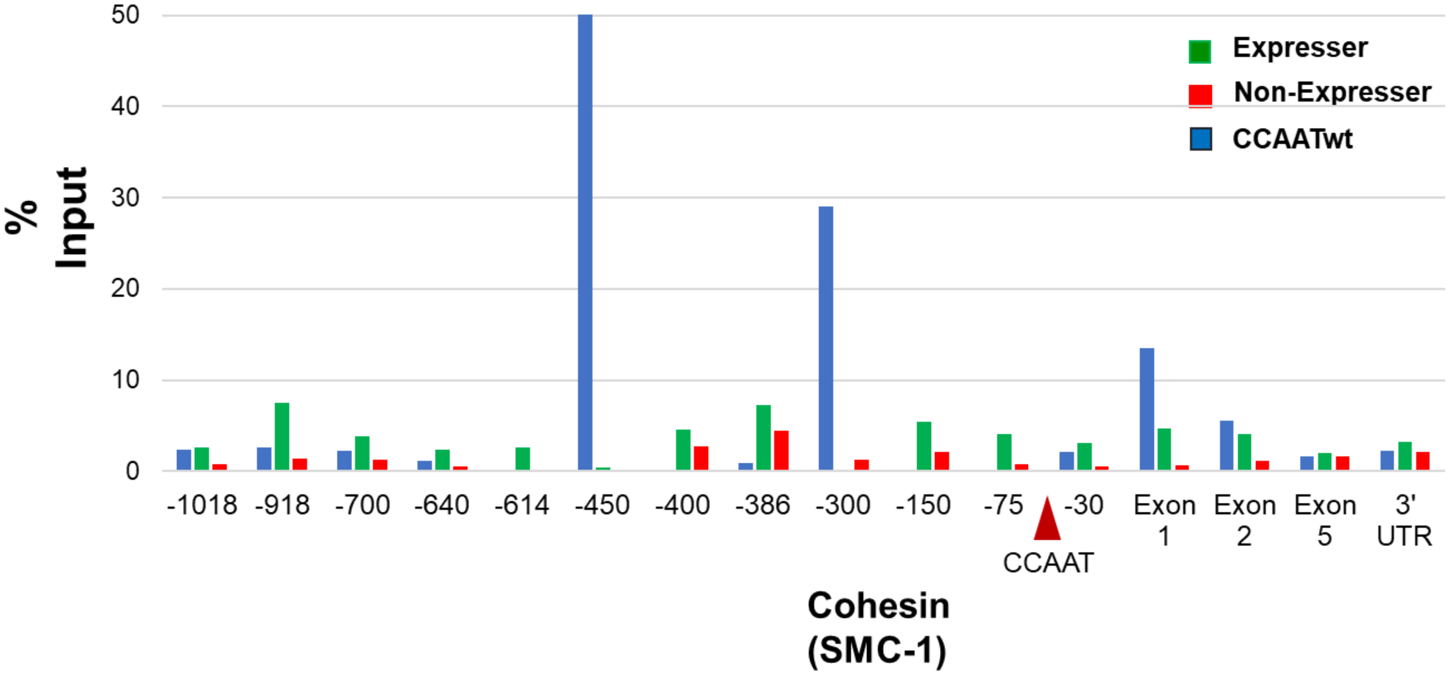


**Fig. 7S: CCAATm mice show similar patterns of cohesin binding**

ChIP analysis of Cohesin (Smc1) binding to chromatin from spleens of CCAATwt, CCAATm expresser and CCAATm non-expresser transgenic strains as % of total Input. The CCAAT mutation location is denoted by the triangle. Note X axis denotes location relative to the TSS and is not to scale. Results are representative of 3 of experiments.
